# Computational and experimental approaches to controlling bacterial microcompartment assembly

**DOI:** 10.1101/2021.01.18.427120

**Authors:** Yaohua Li, Nolan W. Kennedy, Siyu Li, Carolyn E. Mills, Danielle Tullman-Ercek, Monica Olvera de la Cruz

**Author notes:** These authors contributed equally to this work (Yaohua Li, Nolan W. Kennedy).

## Abstract

Bacterial microcompartments compartmentalize the enzymes that aid chemical and energy production in many bacterial species. These protein organelles are found in various bacterial phyla and are postulated to help many of these organisms survive in hostile environments such as the gut of their hosts. Metabolic engineers are interested in repurposing these organelles for non-native functions. Here, we use computational, theoretical and experimental approaches to determine mechanisms that effectively control microcompartment self-assembly. As a model system, we find via multiscale modeling and mutagenesis studies, the interactions responsible for the binding of hexamer-forming proteins propanediol utilization bacterial microcompartments from Salmonella and establish conditions that form various morphologies. We determine how the changes in the microcompartment hexamer protein preferred angles and interaction strengths can modify the assembled morphologies including the naturally occurring polyhedral microcompartment shape, as well as other extended shapes or quasi-closed shapes. We demonstrate experimentally that such altered strengths and angles are achieved via amino acid mutations. A thermodynamic model that agrees with the coarse-grained simulations provides guidelines to design microcompartments. These findings yield insight in controlled protein assembly and provide principles for assembling microcompartments for biochemical or energy applications as nanoreactors.

## Introduction

Compartmentalization of cell components enables a variety of functions and ensures that biochemical processes happen without interfering with one another. Proteins play a crucial role in compartmentalization. Examples of closed compartments formed by protein components include viral capsids, which protect enclosed nucleic acids via robust assembly rules ^1, 2^, and bacterial microcompartments (MCPs), which aid the breakdown of chemicals and energy production that allow bacteria to thrive in various environments ^3^. MCPs, which are polyhedral structures capable of controlling the transport of specific molecules, are found in a large variety of bacterial species ^4–6^. Due to their native ability to enhance chemical transformations within cells, synthetic biologists are working to repurpose MCPs for non-native, industrially relevant purposes ^5, 7^. Therefore, determining and understanding assembly mechanisms of MCPs is a relevant to industry and to life and physical sciences problem.

The envelopes of MCPs are assembled from proteins, much like some viral capsids. In contrast to some capsids, however, these shells are often irregular in shape, contain multiple types of proteins, and enclose specific enzymes as cargo ^8^. Three types of shell protein components form the envelope of 1,2-propanediol utilization (Pdu) MCPs, ethanolamine utilization (Eut) MCPs and many other MCPs ^9^. These three MCP shell components are composed entirely of two different protein domains, the bacterial microcompartment (BMC) domain (which comprises the hexameric and trimeric proteins) and the bacterial microcompartment vertex (BMV) domain (which comprises the pentameric proteins). Six BMC-domain monomers form the flat, six-sided, hexameric structures (BMC-H); three tandem BMC-domain dimers form the flat, six-sided, pseudohexameric trimers (BMC-T), and five BMV-domain monomers form the flat, five-sided, pentameric structures (BMC-P). When hexagonal shape components (BMC-H and BMC-T) assemble with pentagonal components (BMC-P) into closed shells, the topological constraints of the Euler’s polyhedron formula are obeyed ^10^. This formula states that 12 pentagonal components (BMC-P) are needed to form a topologically closed shell, which explains the ubiquitous icosahedral symmetry of spherical shells^11^ and their buckling into icosahedral shells ^12^. However, the closed shells formed by many MCP systems, including the Pdu MCP system, have complex polyhedral geometries ^13–15^ in which shell protein components are present at different ratios ^16, 17^. Polyhedral MCPs are predicted in multicomponent closed shells using elasticity theory ^18, 19^, yet the conditions, including protein size, shape and interactions, to achieve specific structures are unknown. In certain conditions these MCP proteins assemble to produce extended cylindrical structures ^13^. A recent study demonstrated that open “icosahedral cages” can be formed in the absence of BMC-P proteins ^17^. These cages contain open spaces that enable BMC-P binding such that Euler’s polyhedron formula is still obeyed when the shell closes. These studies have shed light on some basic aspects of MCP shell structure. The rules which govern MCP assembly into various morphologies, however, are not well-understood.

While genetic and crystallographic studies have increased understanding of shell formation and MCP architecture, shell protein mutation studies have also provided information on specific shell protein functionality ^3^ including structure and transport ^20^. Recent mutation studies that alter the charge of pore amino acids in Pdu MCPs, for example, showed how pore mutations affect growth on 1,2-propanediol^7^. BMC-H proteins are found at high abundance in the MCP shell ^21^ and play a crucial role in MCP assembly^22^, raising the importance of inter-hexamer interactions in the molecular layer presumed to comprise the facets of the shell.

Here, we develop a multi-scale computational approach and a thermodynamic model combined with mutagenesis studies and experimental observations to analyze the assembly of native and mutated Pdu MCP shell proteins. As an experimental model system we study the PduA BMC-H protein of *Salmonella enterica* serovar Typhimurium LT2 ^23^. By computationally studying specific interactions between PduA hexamers, we find the conditions for assembly into closed polyhedral or extended shapes and examine a specific mutation site on these hexamers that affects the assembly of MCPs into different morphologies. We use all-atom (AA) molecular dynamics (MD) simulations to determine the interaction energy and equilibrium bending and twisting angles of native and mutated hexamers and predict what mutations can lead to self-assembly of MCPs or extended shapes. Experimental results with mutated PduA hexamers qualitatively corroborate the simulation predictions. Our study reveals that electrostatic and hydrogen bonding interactions between the arginine and valine residues on the edge of PduA are key to the self-assembly of PduA hexamers and MCP formation, as the majority of the mutations to this residue negatively impact MCP formation ^24^. We use the structural information and interaction energy from AA MD simulations to construct a coarse-grained (CG) model for PduA self-assembly. Moreover, we determine what assembled geometries are accessible given different stoichiometric ratio of the 3 major MCP component proteins. Finally, we use this CG framework in combination with a theoretical thermodynamic model to explore how modulating BMC-H/BMC-H and BMC-H/BMC-P interactions can permit access to an array of assembled morphologies. Simulation and theoretical findings agree with *in vivo* experiments of MCP encapsulating green florescent proteins (GFP). These findings will help guide future studies that seek to repurpose MCPs with specific morphologies and shed light onto the rules governing MCP assembly in general.

## Results

### Molecular Dynamic Simulations

We first investigate the interaction between two native BMC-H subunits using AA MD simulations of the PduA crystal structure from *S. enterica* LT2 (PDB id: 3ngk)^25^. The two PduA hexamers bind into a highly stable dimer (see Fig. 1A and SI video 1) as a result of complementary shape and hydrogen bonding (see Table 1), in which the arginine at the 79^th^ position (ARG79) binds to the backbone carbonyl oxygen of VAL25. The side chain of ARG79 fits in the pocket formed by the residues ASN29, VAL25, HIS81 and LYS26 (Fig. 1A inset). This result corresponds to previous studies indicating ARG79 plays a role in inter-hexamer binding ^24, 26–28^. To better quantify the relative orientation of the two hexamers, we calculated the angle between the norm of the hexagonal planes (Fig. 1B). The angle is projected onto the y-z plane and x-z plane to decompose into a bending angle *θ*_*b*_ = 10 ± 4° (error bars are standard deviations) and a twisting angle *θ*_*t*_ =5± 3° at equilibrium (Fig. 1C), with a mean total angle *θ* = 11 ± 4°. (A histogram of the bending and twisting angles are shown in SI Fig S1). This calculation suggests that in solution PduA hexamer complex mostly adopts a near coplanar configuration, corresponding to the flat facets observed in polyhedral MCP structures ^27^; there is another orientation with the hexamers at a 30° angle relative to each other ^27^ (see SI Fig. S2). The latter bent configuration is likely due to collective interactions among other proteins and scaffold, as shown in our coarse-grained model described below, and therefore, it is not observed in simulations of two proteins.

**Figure 1.**
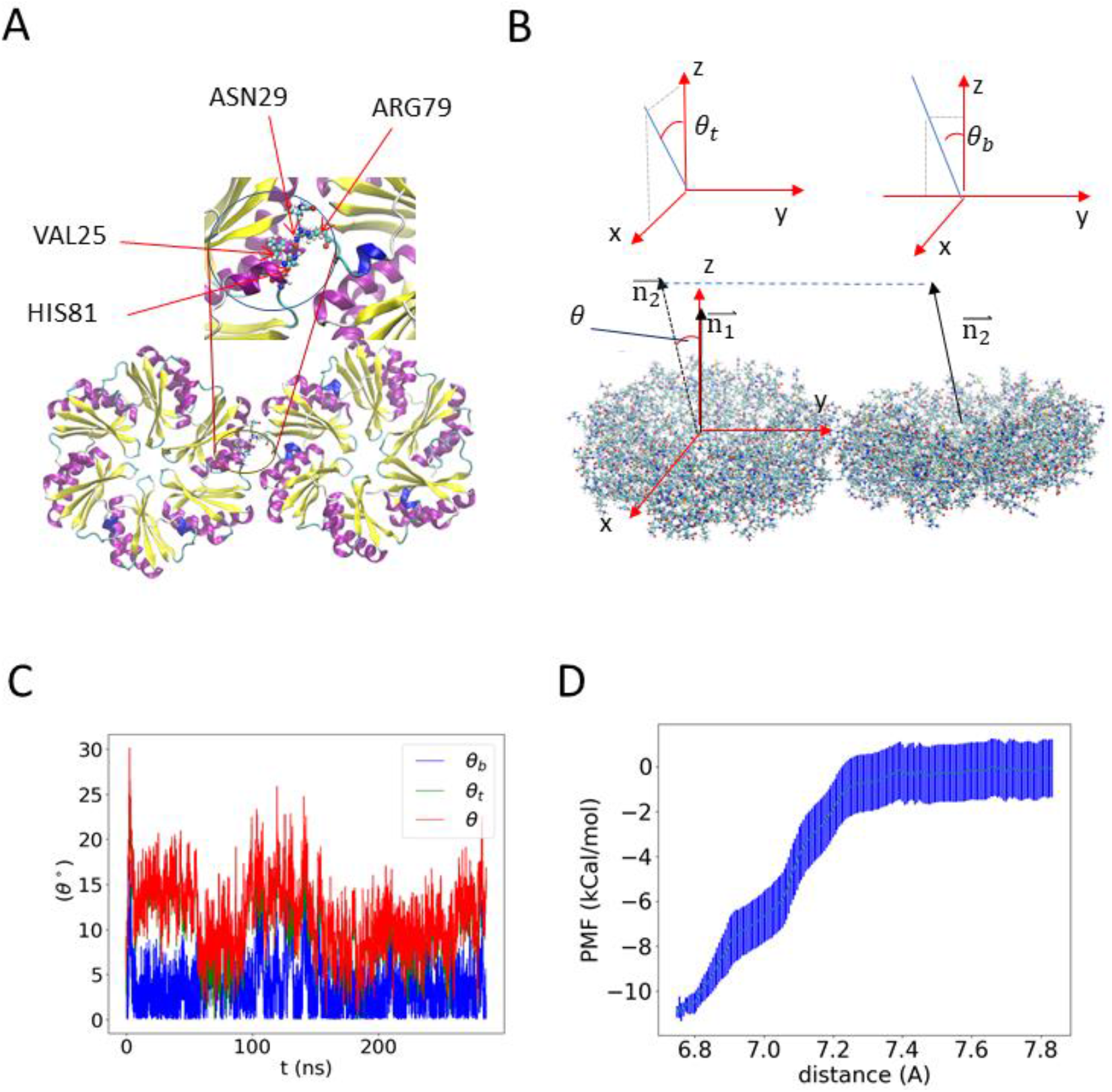
All atom (AA) molecular dynamics (MD) simulations results of a pair of PduA hexamers. (A) The equilibrium configuration. The arginine residue on each edge sticks out to form hydrogen bond with the backbone of valine in the neighboring PduA hexamer. The two arginine residues are highlighted by increased bead size. Water and ions are hidden for visual clarity. (B) Illustration of the reference frame and angle between the two PduA hexamers. The angle *θ* is decomposed into twisting angle *θ*_*t*_ and bending angle *θ*_*b*_ by projecting the normal of the neighboring protein (**n_2_**) onto the XZ plane and the YZ plane, respectively (see SI Fig. S1 for a different representation) (C) The MD simulation values of *θ*, *θ*_*t*_ and *θ*_*b*_. (D) Potential of mean force between two PduA hexamers shows an attractive binding energy of about 11 kCal/mol.

**Table 1.**
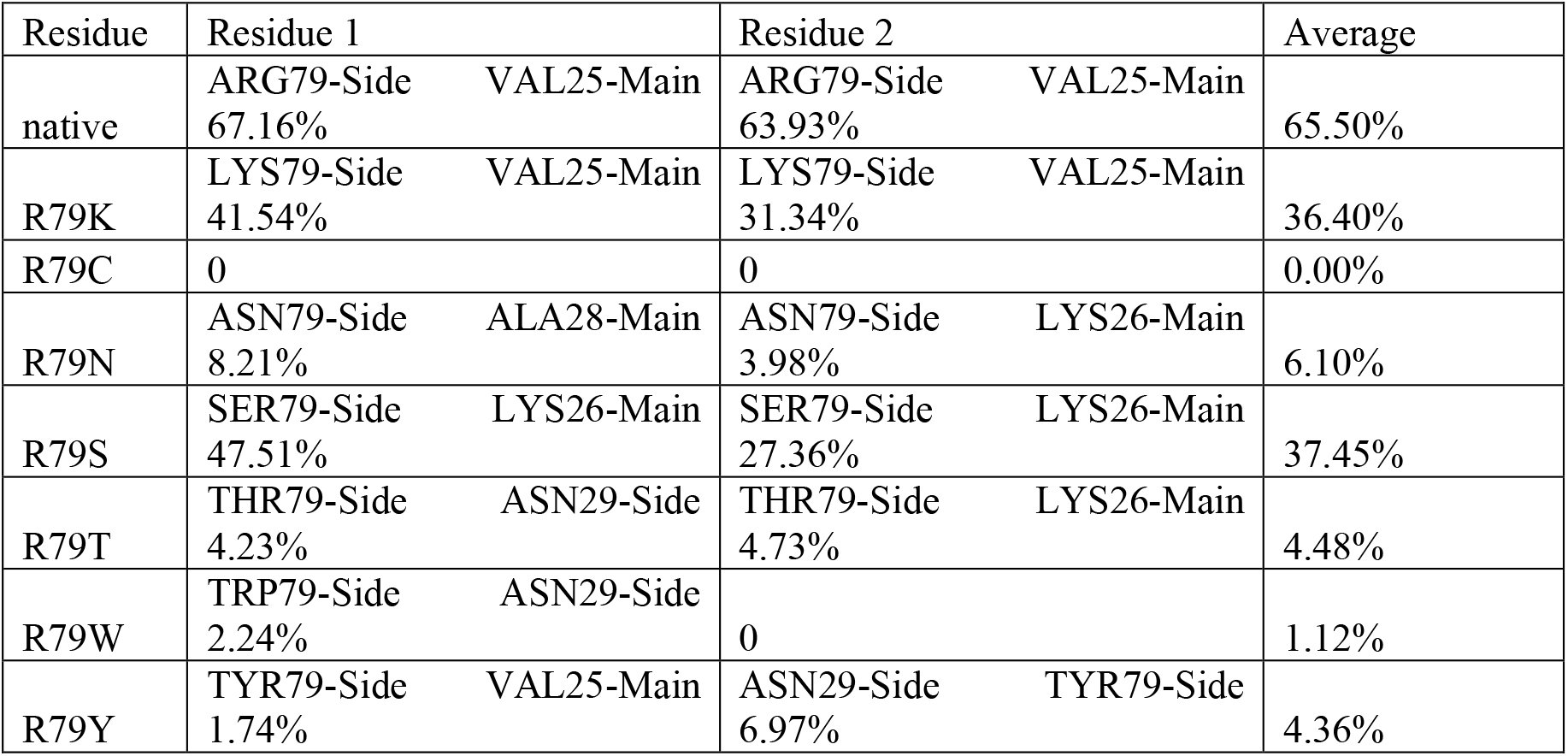
Hydrogen bond analysis of selected PduA mutants (listed as a percentage of time during the simulation)

We study the potential of mean force (PMF) of two PduA hexamers using umbrella sampling MD simulations (Fig. 1 D). The binding energy (*ε*_*hh,AA*_) is estimated to be 11 ± 2 kCal/mol. This binding energy falls in the range of reported values of hydrogen bonds ^29^, further indicating that hydrogen bonds are a major contributor to PduA binding. The calculated binding energy and bending and twisting angles from atomistic simulations provide semi-quantitative guidance for building larger scale models.

Since AA simulations cannot include many proteins and cover the time scales required to assemble the proteins into different MCPs morphologies, we use CG modeling to study the assembly of MCPs at larger time and length scales. CG simulations have been used to follow the assembly of simplified protein models into MCPs or viral capsids ^30, 31^. MD simulations of one shell protein predict the stable formation of small icosahedral capsids^32^. CG models have also elucidated the initial faceting process^33^ and the formation of a complete MCP ^30^. However, these models have not investigated how interactions between multiple shell components result in various MCP morphologies such as cylinders, and the molecular origin of spontaneous curvature has not been explained. We hypothesize that the stoichiometric ratio of different components and the interaction strength between them control the morphology of assembly products ^9, 14, 34^. We construct a CG model using the approximate shapes and charge distributions of the aforementioned PduA structure (PDB: 3ngk^25^) for the BMC-H model, the PduB homolog from *Lactobacillus reuteri* (PDB: 4fay^35^) for the BMC-T model, and the PduN homolog GrpN from *Rhodospirillium rubrum* (PDB: 4i7a^36^) for the BMC-P model. We validate this model by comparing assembled morphologies against known experimental observations of various MCP and cylinder shapes^9, 13, 14^.

The components of our CG model are shown in Fig. 2A. In the CG simulations the charges of the BMC-H, BMC-T and BMC-P proteins are computed from the AA structures (see SI Fig. S3), the mechanical properties such as the spontaneous curvature of BMC-H assemblies result from the inclined edge shape; that is, the inclination angle *θ*_*i*_ ~ 25 ° obtained from the PDB (see CG simulations results using other inclination angles in Figs S4-S5 in SI). We first explore the various geometries with different stoichiometric ratio of BMC-H and BMC-P. We find icosahedral shells assembled with BMC-H: BMC-P = 5: 3, which resemble the shape of *in vitro* MCPs from *Haliangium ochraceum*^13, 27^ (Fig. 2B). When we reduce the content of BMC-P to BMC-H: BMC-P = 6: 1, the addition of hexameric proteins allows the shell to assemble into the irregular shapes of Pdu MCPs reported in the literature^9, 14, 34^, illustrating how varying shell protein content can control morphology of MCP structures formed. When a small portion of BMC-H is replaced by BMC-T, similar structures are found (Figs. 2C and 2D). However, when the content of BMC-T further increases, the asphericity increases (see SI Fig. S6). A detailed structural analysis of assembled MCPs is available in Table S1. The impact of BMC-T on the assembly can be understood by the difference in bending angle and rigidity of the BMC-H and BMC-T (a bending angle analysis is given in the SI). We note that the BMC-H: BMC-P ratio used in these simulations is lower than that typically observed for Pdu MCPs *in vivo* and the MCPs simulated are about 20-40 nm in diameter, compared to typical reported Pdu MCP diameters of 40-600 nm ^15, 34, 37, 38^. These smaller MCPs equilibrate faster, allowing more efficient exploration of parameter space while still qualitatively reproducing the characteristic polyhedral shapes of MCPs observed in experiments^13, 14^. Simulations with BMC-H: BMC-P ratios of 8:1 produced bigger, more aspherical shells (SI Fig. S7).

**Figure 2.**
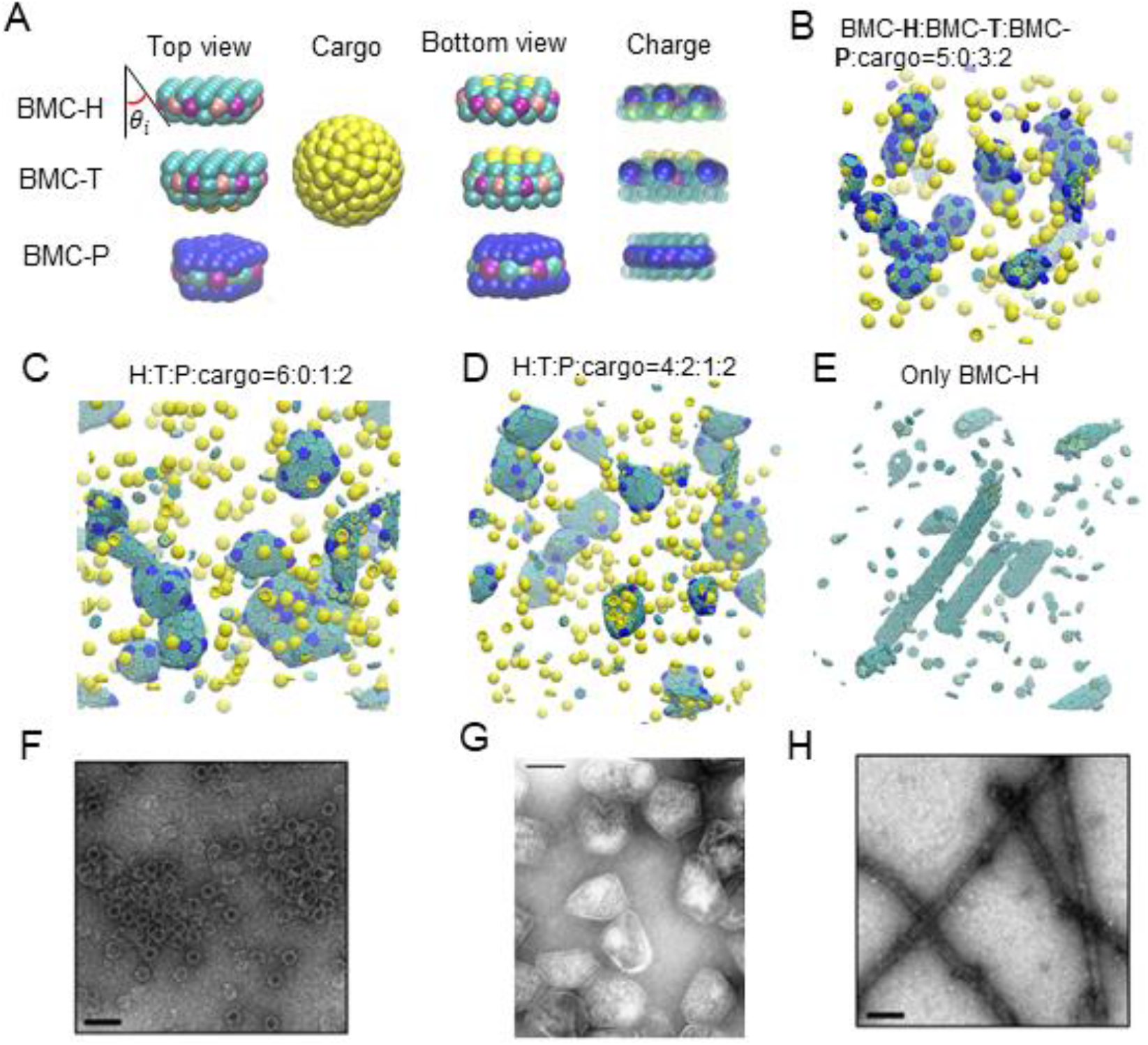
Coarse-grained (CG) model and MD simulations results. (A) Illustration of the CG model. The sides of BMC-H and BMC-T are inclined at an angle *θ*_*i*_ = 25° according to their pdb structures (PDB id 3ngk ^25^ and 4fay ^35^). The green beads interact via excluded volume (changed to blue color for pentamers), the purple and pink beads are short-range attractive sites representing the arginine hydrogen bonds. The sphere of yellow beads is the cargo and the yellow beads on the BMC-H and BMC-T proteins are the residues that bind to the cargo. To show the negative (blue) and positive (green) charged sites clearly, all the non-charged sites are shown as semi-transparent and in smaller size on the right column. (B)~(E) Snapshots of CG simulations (the hexamers, pentamers and cargo are in green, blue and yellow, respectively): a red dot is marked on the center of BMC-T to distinguish them from BMC-H. (B) A system without BMC-T, with a ratio BMC-H:BMC-P:cargo=5:3:2 forms T=3 icosahedral shells resembling in vitro electron micrographs of compartments from *Haliangium ochraceum* (F) which form T=9 shells. (C)~(D) Increasing the number of BMC-H or adding BMC-T can enable assembly into polyhedral shapes that resemble the shape of purified MCPs shown in (F). (E) BMC-H proteins alone form cylinders, reproducing observations of in vitro BMC-H assembly shown in (H). Detailed model parameters for these simulation results are provided in SI Table S1. (G) is reprinted with permission from C. Fan, et al. *PNAS* **2010**, 107, 7509-7514 ^14^, (F) and (H) are reprinted with permission from A. R. Hagen, et al. *Nano Letters* **2018**, 18, 7030-7037 ^13^.

Long cylinders are obtained when only BMC-H proteins are in the assembly, as shown in Fig. 2E. The shape is consistent with both *in vivo* and *ex vivo* experiments ^26, 39^, in which overexpressing PduA proteins resulted in the assembly of long cylindrical tubes, and *in vitro* experiments in which the BMC-H from *H. ochraceum* spontaneously assembled into tubes in the absence of other shell proteins^13^. The ability to reproduce vastly different shapes (cylinders and polyhedra) suggests that this simple CG model captures the key interactions of Pdu MCP proteins.

Once our CG model has been validated by comparing the results with experimentally observed morphologies at different stoichiometry, we explore how the docking angle of BMC-H proteins impacts MCP assembly. AA MD simulations reveal (Table2) that the docking angle of BMC-H can be tuned by mutating the arginine at the binding site. To this end, we use a combination of AA MD simulations, mutation experiments, and CG modeling to determine the role of hexamer-hexamer interactions on BMC-H assembly. AA MD simulations, the PduA conservation score (SI Fig. S8) and prior work in the field^8, 24, 26, 27^ indicate that ARG79 is a key residue for controlling PduA interactions. Thus, we hypothesized that mutations to the arginine residue at the 79^th^ position (ARG79) of PduA could impact PduA, and, subsequently, MCP assembly. We perform AA simulations of the hexamer-hexamer interface on fourteen PduA mutants. Of these, eleven mutants retain some degree of hexamer-hexamer interfacial contact for the duration of the simulations, and their bending and twisting angles are shown in Fig. 3A-C. Three mutants result in hexamers that entirely dissociate from each other, completely eliminating interfacial contacts (Table 2 and Fig. S9 in the SI provide further details). The hexamer-hexamer dissociation arises when the arginine is replaced by a negatively charged residue or hydrophilic residue with low isoelectric point (aspartic acid, glutamic acid and glutamine), indicating that Coulombic interactions play an important role in stabilizing the hexamer-hexamer interface in PduA. The alpha-helix of the opposite hexamer has a negative partial charge, which is thought to complement the native, positively charged arginine residue^28^. Therefore, mutating the arginine to the opposite charge result in total dissociation of the PduA.

**Figure 3.**
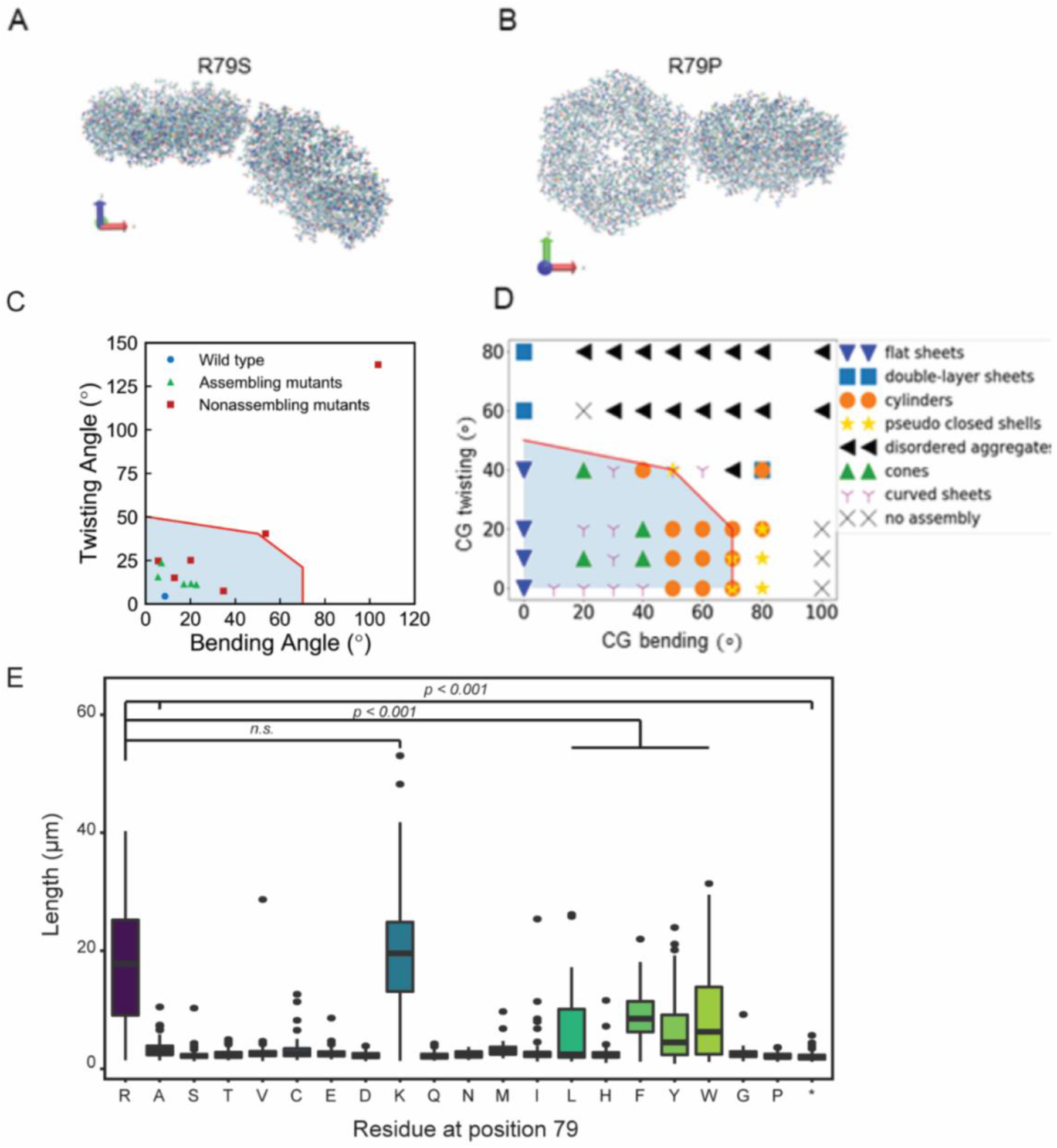
PduA mutant orientation studies. An example of a mutation with large bending angle is shown in (A) and large twisting angle in (B). (C) The twisting and bending angle calculated for 11 R79 mutation from AA simulations (corresponding to the data in Table 2). The PMF for 3 mutations and WT are calculated and shown in Table 2. The red line indicates the critical angle to form extended structures as predicted by CG simulations. Triangle and square markers indicate whether the mutants self-assemble in experiments (E and SI Fig. S10) (D) The morphologies formed in CG simulation of 256 PduA proteins with given bending and twisting angles. The shaded area below the solid red lines in (C) and (D) indicate the system can form extended structures, which we predict to cause long chains of cells to form in experiments. At bending angles of 70 and 80, the pseudo closed shells (yellow stars) resemble quasi-icosahedra of T=1 in that each hexamer has 5 neighbors (SI Figure S11). (E) Distribution of cell length populations for each PduA variant. R79R (WT) and R79K were significantly longer than other PduA variants (p < 0.001, t-test).

**Table 2.**
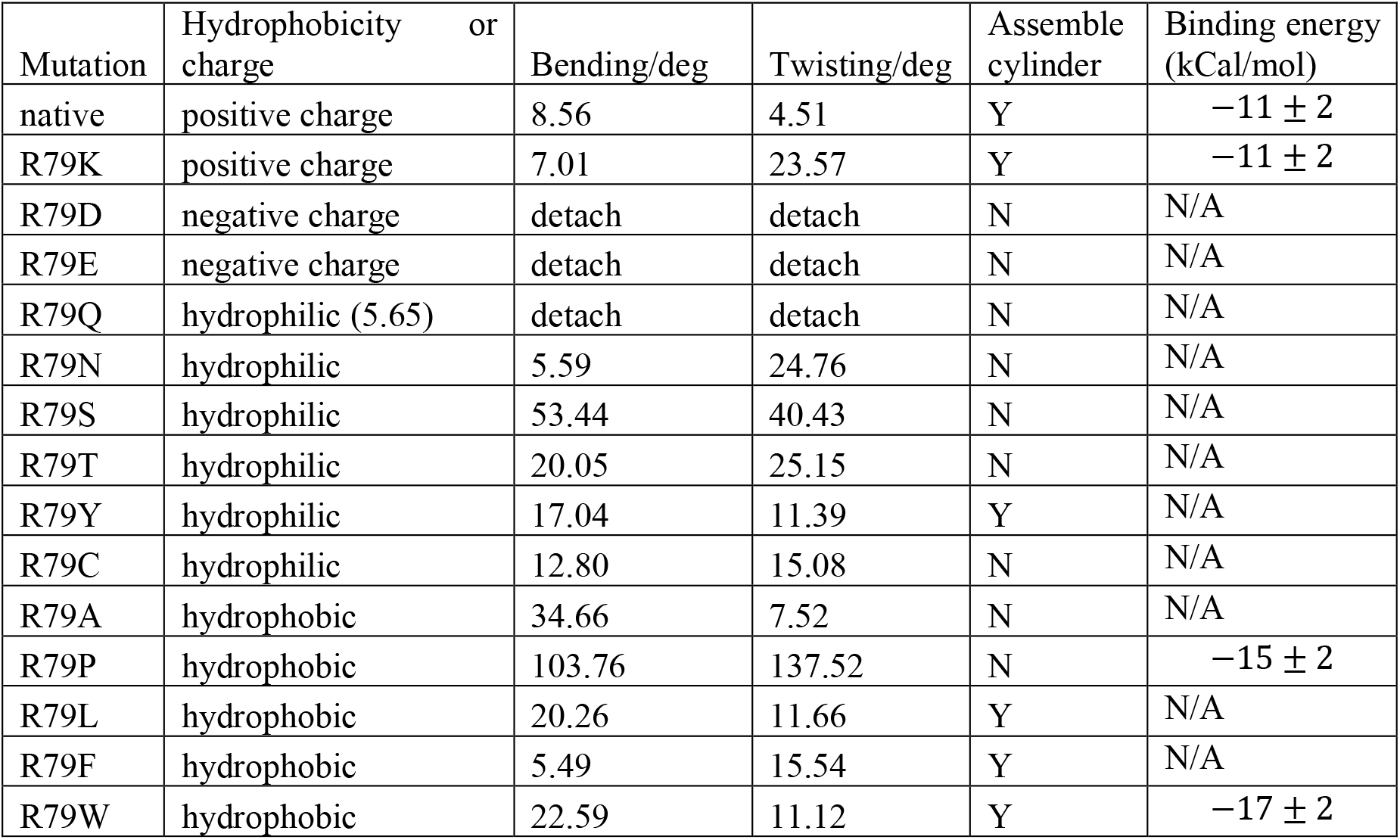
Summary of all atom molecular dynamics simulation results. (In the binding energy column, negative values correspond to attractive energy)

| Mutation | Hydrophobicity or charge | Bending/deg | Twisting/deg | Assemble cylinder | Binding energy (kCal/mol) |
| --- | --- | --- | --- | --- | --- |
| native | positive charge | 8.56 | 4.51 | Y | $-11 \pm 2$ |
| R79K | positive charge | 7.01 | 23.57 | Y | $-11 \pm 2$ |
| R79D | negative charge | detach | detach | N | N/A |
| R79E | negative charge | detach | detach | N | N/A |
| R79Q | hydrophilic (5.65) | detach | detach | N | N/A |
| R79N | hydrophilic | 5.59 | 24.76 | N | N/A |
| R79S | hydrophilic | 53.44 | 40.43 | N | N/A |
| R79T | hydrophilic | 20.05 | 25.15 | N | N/A |
| R79Y | hydrophilic | 17.04 | 11.39 | Y | N/A |
| R79C | hydrophilic | 12.80 | 15.08 | N | N/A |
| R79A | hydrophobic | 34.66 | 7.52 | N | N/A |
| R79P | hydrophobic | 103.76 | 137.52 | N | $-15 \pm 2$ |
| R79L | hydrophobic | 20.26 | 11.66 | Y | N/A |
| R79F | hydrophobic | 5.49 | 15.54 | Y | N/A |
| R79W | hydrophobic | 22.59 | 11.12 | Y | $-17 \pm 2$ |

### Mutagenesis Experiments Comparison with Simulations of Hexamers Assembly

We turned to an experimental approach to assess the assembly of PduA mutants and validate the above described AA MD simulations. When PduA is overexpressed in *E. coli* cells it naturally assembles into long cylinders that span the length of the cell body. The assembled protein structures form bundles within the cell cytoplasm, preventing cell division and leading to extended chains of cells^22^ (SI Fig. S10). This allows for cell length to be used as a proxy for protein assembly and allows us to rapidly assess the assembly state of different PduA mutants. We hypothesized that mutants which formed stable dimers and had limited bending and twisting angles in AA MD simulations would be more likely to lead to the linked cell phenotype, indicative of protein self-assembly into cylinders. Indeed, the two PduA variants predicted to form hydrogen bonds (the native ARG79 and the R79K mutant, where “R79K” indicates that the arginine (R) at 79^th^ position in the protein sequence is mutated to a lysine (K)), have among the lowest bending and twisting angles in simulations (Fig. 3C, Table 2), and overexpression of both proteins leads to highly-elongated cells (Fig. 3E, SI Fig. S10) compared to the negative control mutants (R79A and R79*, where “*” indicates a stop codon) (p < 0.0001). We also verified that the expression level of various PduA mutants did not correlate strongly with cell length, indicating that differences in assembly state are not likely to be due to differences in expression (SI Fig. S11). These results indicate successful assembly of WT PduA and PduA-R79K into cylinders. Surprisingly, we found that substitution of ARG79 with large hydrophobic residues (R79L) or aromatic residues (R79F, R79Y, R79W) also enabled some amount of assembly, as indicated by the presence of linked cells during overexpression (Fig. 3E, SI Fig. S10), although to a lesser degree than the native ARG79 (p < 0.001). While initially surprising, this assembly is predicted by the low bending and twisting angles calculated for these mutants from our simulations (Fig. 3C). Free energy calculations for the binding energy of select mutants are also provided (R79K, R79W and R79P; see SI Table 2), but these do not correlate as well with cell length as bending and twisting angles—while hydrophobic residues tryptophan (R79W) and proline (R79P) have similar binding energies, experiments indicate that they do not exhibit similar self-assembly behavior. Overall, two residues that form hydrogen bonds (R79R and R79K) and aromatic residues (R79W, R79F and R79Y) both confer small bending and twisting angles between hexamers. Together, these results imply that bending and twisting angles as determined by our AA MD simulations can provide insight into the assembly of PduA.

The combination of experimental and AA simulation results described above suggests that PduA assembly is largely influenced by the inclination angle *θ*_*i*_ and twisting angle *θ*_*t*_ between hexamers. To understand what assembled morphologies may be accessed by these different PduA mutants, we performed CG MD simulations on PduA alone (no other proteins and no cargo) using the BMC-H model described above, and documented how preferred inclination and twisting angle between hexamers impacts the morphology of PduA-only structures formed (Fig. 3D). From the inclined geometry assumed in the CG model, the resulting bending angle between two hexamer planes is approximately 2*θ*_*i*_ (SI Fig. S4A). The CG model predicts that at small *θ*_*i*_, the mutated PduA assemble into extended structures (flat/curve sheet, cylinder/cone) with low curvature, whereas at large *θ*_*i*_, the hexamers form structureless aggregates or non-extended pseudo-closed shells with high curvature (See SI Fig. S12 for snapshots of assembled morphologies). The flat or curved sheets are expected to roll into cylinders or cones in the presence of cargo and cell confinements. The predicted critical angles for self-assembly into extended structures from CG simulations are given by the red line in Figs. 3C and 3D. We note that several non-assembling mutants have bending and twisting angles (calculated by AA simulations) below this line. This quantitative discrepancy is because AA simulations have detailed interatomic interactions omitted in the CG model that overestimate the CG critical bending and twisting angle. That is, agreement with the *in vivo* experiments is found if we take into account this shift by moving red line to the lower left. Taken together, the AA and CG simulations indicate that pure PduA hexamers that form interfaces with small bending and twisting angles can assemble into extended structures. Mutations with big angles, despite having comparable attractive interaction strength, tend to form pseudo closed shells, unstructured aggregates and stacked layers (see Fig. S12 in SI).

### Multicomponent Microcompartment Assembly: Thermodynamic Model Comparison with Coarse-Grained Simulations and Experiments

Extending our exploration of how shell protein interactions impact MCP assembly, we next used CG MD simulations and thermodynamic modeling to explore how hexamer-hexamer and pentamer-hexamer interaction strengths impact assembled morphologies. We hypothesized that modulating these interaction strength ratios would allow us to tune the morphologies of MCPs formed, enabling downstream engineering efforts. The thermodynamic model is constructed by numerically minimizing a free energy that includes protein-protein binding, protein-cargo interaction, elastic penalty of shell bending and chemical potentials (see Materials and Methods), using parameters consistent with the CG model described in the previous section. Based on the hexamer-hexamer binding energies calculated in our R79 mutation simulations, we specified a range of accessible BMC-H/BMC-H (*ε*_*hh*_) and BMC-H/BMC-P (*ε*_*ph*_) interaction energies. We used a protein ratio of BMC-H:BMC-T:BMC-P:cargo = 4:2:1:2 for CG simulations (Fig. 4A). For simplicity, the interaction parameters for BMC-T/BMC-T and BMC-T/BMC-H were set equal to the BMC-H/BMC-H interaction energy, because PduB trimers have conserved arginines in positions analogous to the arginines on PduA hexamers, and these are thought to be critical for binding^27^ (in the SI Fig. S6 D, we discuss the effect of changing these values). The data shown in Fig. 4A are for a BMC-H and BMC-T inclination angle of *θ*_*i*_ = 25°; however, data taken for an inclination angle of 15° produces qualitatively similar morphologies and trends (SI Fig. S4B), suggesting that the observed trends hold for a variety of inclination angles.

**Figure 4.**
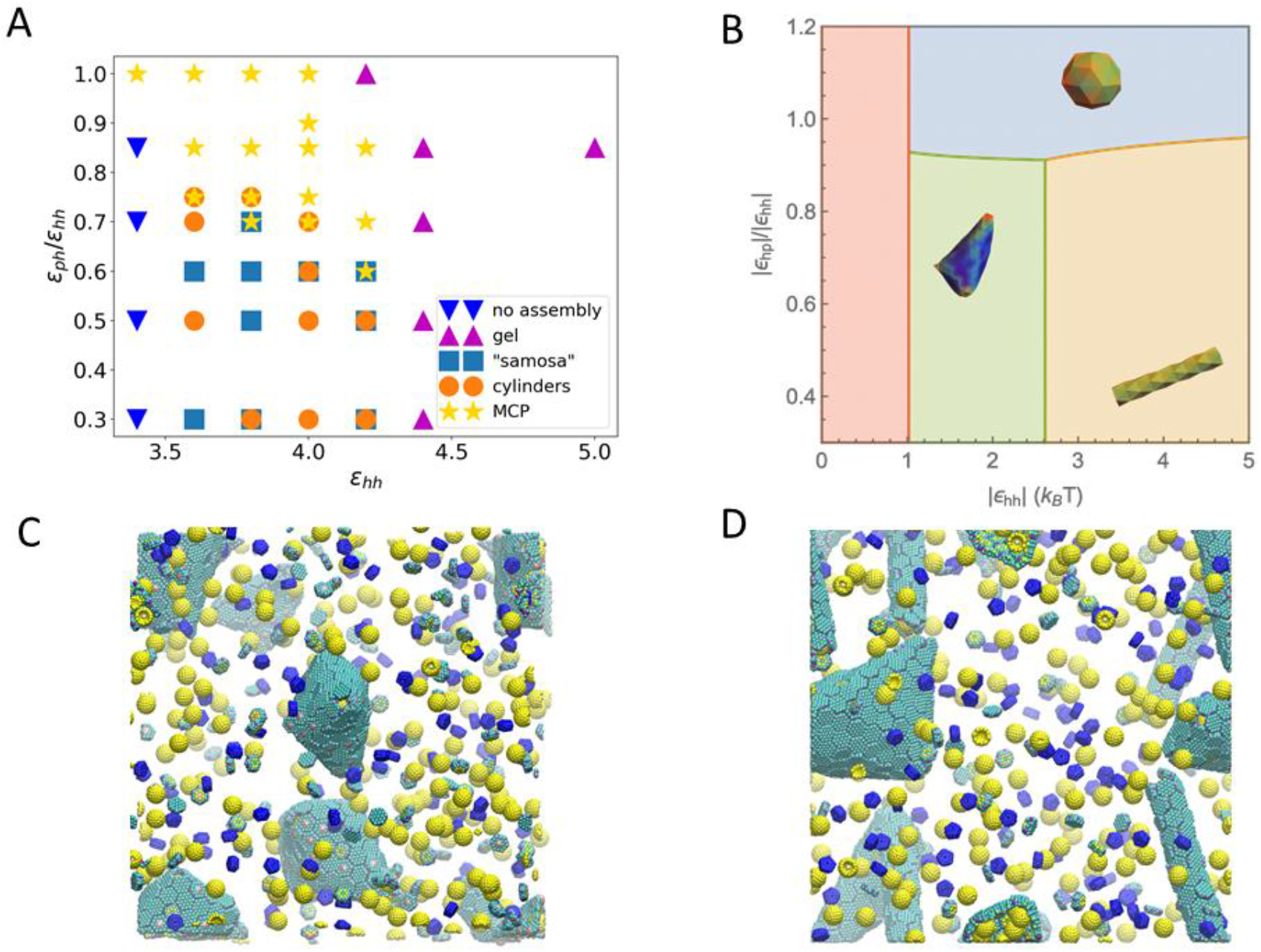
(A) Phase diagram of assembled shapes from CG MD simulations. The axes are the *ε* parameters defined in Eqn.1. The simulation box is constructed from a unit of 4 BMC-H, 2 BMC-T, 2 enzymes, and 1 BMC-P, replicated 5 times in x, y and z directions. (B) Phase diagram from thermodynamics analysis (see parameters used in Table S5 in SI). When the hexamers binding energy is weak, there is no assembly (red region). With stronger hexamer-hexamer interaction *∊*_*hh*_, MCP, cylindrical and samosa shaped shells are formed, corresponding to the blue, yellow and green regions, respectively. (C) and (D) are snapshots of CG simulations (the hexamers, pentamers and cargo are in green, blue and yellow, respectively). (C) The “samosa” shape (*ε*_*hh*_ = 3.8, *ε*_*ph*_ = 1.9) is a quasi-closed surface without BMC-P proteins. They are different from MCPs in that they have no pentamers, and the vertices are sharp cones with a hole or defect at the tip. An example of a 4-fold defect is shown in the center. (D) Co-existence of cylinders and “samosas” at (*ε*_*hh*_ = 3.8, *ε*_*ph*_ = 2.28).

The different morphologies predicted by the CG simulation and by the thermodynamic theory as a function of pentamer-hexamer and hexamer-hexamer interactions are shown Figs. 4A and 4B, respectively. The phase diagram constructed using CG MD simulations shows that closed MCP structures form when 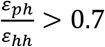 and *ε*_*hh*_ is between 3.6 and 4.2. This range of energy values corresponds to a hexamer-hexamer attraction energy between 4 and 6 kCal/mol, obtained using Eq.1, which does not include the repulsion caused by screened electrostatics (adding this screened Coulomb energy contribution, the overall attractive energy is between 3.6 and 5.6 kCal/mol). When hexamer-hexamer interaction strength is increased beyond this MCP-forming range (*ε*_*hh*_ > 4.4), the proteins bind irreversibly and assemble into gel-like structures. When hexamer-hexamer attractions are weakened (*ε*_*hh*_< 3.4), no assembled morphologies are observed. When *ε*_*ph*_ is small compared to *ε*_*hh*_ 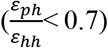, assembled structures do not contain pentamers. These structures include cylinders and structures resembling “samosas” with four-fold coordinated defective holes at the corners (Fig. 4C). We note that these samosa-like structures are also observed when inextensible sheets are folded to form quasi-closed surface, for example in macroscopic inelastic membranes ^40^.

The thermodynamic theory (described in Materials and Methods) predicts changes in morphology with changing pentamer-hexamer (*∊*_*ph*_) and hexamer-hexamer (*∊*_*hh*_) interaction energies. By comparing the formation energy of each tested morphology, we obtain the phase diagram shown in Fig. 4B. In the region of *∊*_*hh*_ < 1.0*k*_*B*_*T* (in red), the nucleation barrier is large, and, as a result, no shells could be formed. At a moderate value of *∊*_*hh*_, the shells can assemble into multiple shapes depending on the value of *∊*_*ph*_. When *∊*_*ph*_ is sufficiently high, the shell assembles into a small closed shell (MCP) (blue region). In the lower pentamer-hexamer interaction region, hexamers aggregate without pentamers, and in such case, cargo is required to provide the curvature to close the shell. The competition between cargo-hexamer interaction and the bending energy decide whether a cylindrical shape or a “samosa” shape is assembled. While the cylindrical shells can encapsulate dense cargo at a considerable bending energy cost, the “samosa” shell can encapsulate loose cargo at a smaller bending energy penalty. These results are in qualitative agreement with the CG simulations in Fig. 4A, except that the CG simulations for large *ε*_*hh*_ values produce gels of connected sheets trapped in a local minimum. The lack of quantitative agreement can be explained by the fact that the thermodynamic model assumes equilibrium and constant reservoir concentrations during the assembly.

Many of the morphologies observed in our CG simulations (Fig. 4) have been observed in the literature for different MCP systems assembled under various conditions (Fig. 5) ^14, 26, 36, 39^, supporting our hypothesis that shell protein interaction strength plays an important role in determining MCP morphology. To provide experimental support for the observed effects of changing hexamer-hexamer binding energy on MCP assembly in our models, we used a green fluorescent protein (GFP) encapsulation assay to probe MCP formation in strains expressing the PduA mutants characterized above. In this assay, GFP is fused to an N-terminal signal sequence that is sufficient for targeting GFP to the lumen of MCPs ^14, 41^. This results in bright, fluorescent puncta in the cell cytoplasm if MCPs are present (Fig. 6 A). However, if MCPs do not form properly, fluorescence will be observed instead at one or both poles of the cells; these are termed polar bodies (Fig. 6A). In this way, it is possible to determine if a strain is capable of forming MCPs.

**Figure 5.**
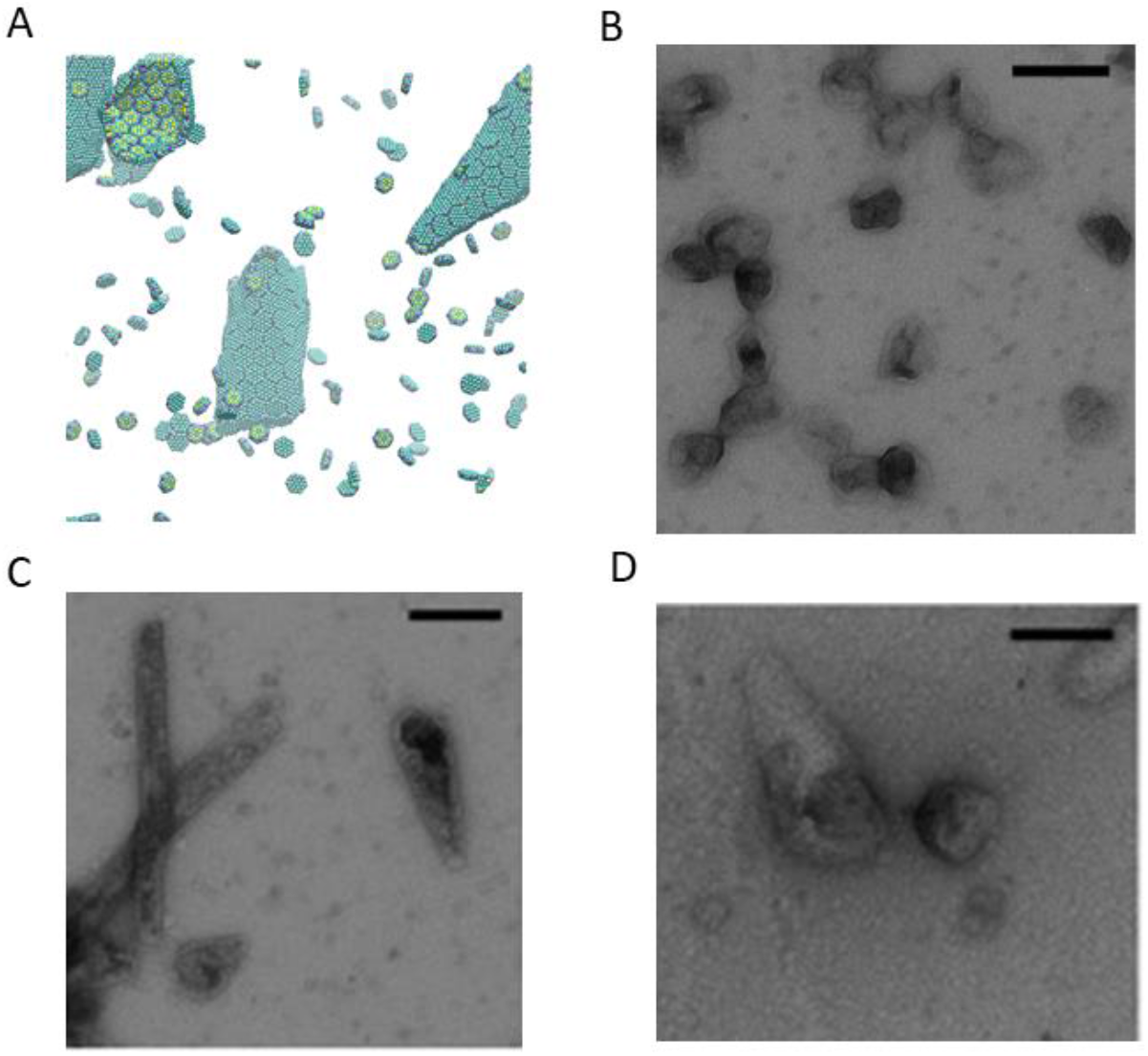
Additional morphologies formed in the CG model and comparisons to morphologies observed by TEM in experiments. (A) With a slightly smaller tilting angle, a system of only BMC-H can form cones similar to the ones observed in (C) and (D). (B) native MCP shells have similar shape compared to simulated MCPs with BMC-H, BMC-T, and BMC-P in Fig. 2C and D. When BMC-P has weaker interactions (ε_hh_ = 3.8, ε_ph_ = 2.28) as shown in Fig. 4 D the proteins form coexisting cylinders and “samosas”, similar to those observed in (C) and (D). (B)~(D) are reprinted with permission from T. M. Nichols, et al. *Biochemical Eng. J.* **2020**, *156*, 107496 ^16^.

**Figure 6.**
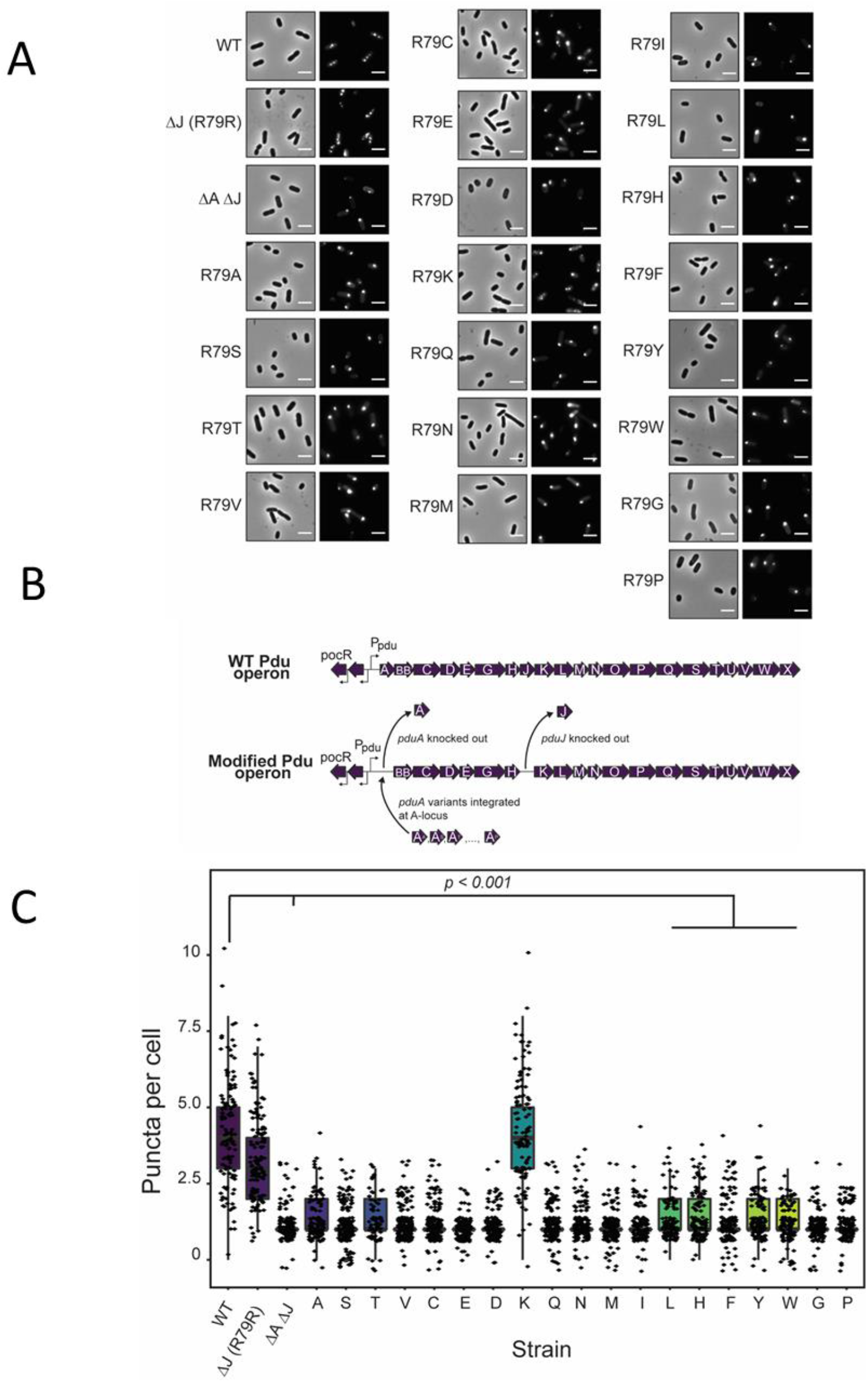
(A) GFP encapsulation assay for PduA variant strains. Phase contrast microscopy (left) and fluorescence (right) micrographs of modified Salmonella strains. GFP-containing MCPs appear as bright puncta in the cell cytoplasm (see WT image), while malformed MCPs appear as polar bodies (see ΔA ΔJ image). Scale bar = 3 μm. (B) Pdu operon modification strategy. Schematic representation of strains with a modified Pdu operon. WT PduA is replaced with PduA variants and PduJ is knocked out in all variant strains. (C) Population distribution of puncta (MCPs) per cell for each strain. ΔJ and ΔA::R79K ΔJ had significantly more puncta per cell than other variant strains and the negative control ΔA ΔJ strain (p < 0.001, t-test).

To this end, we constructed 19 mutant strains of *Salmonella enterica* serovar Typhimurium LT2 to test the effect of altering PduA assembly on overall MCP assembly. Alterations were made to the *pdu* operon as shown in Fig 6B. Because PduJ is capable of overcoming loss of PduA function ^22^, we have knocked out the *pduJ* open reading frame from these strains. Thus, the WT and ΔJ strains serve as positive controls, as these both contain functional, WT PduA. Note that fluorescent puncta are visible throughout the cytoplasm of these strains, as expected when MCPs form (Fig 6A). The ΔA ΔJ strain, in which both the *pduA* and *pduJ* open reading frames have been knocked out, serves as a negative control. Only polar bodies are observed in this strain, indicating improper MCP assembly (Fig. 6A). In the experimental strains, a point mutant of PduA is encoded at the *pduA* locus (Fig 6A) and puncta were counted in each cell to determine the effect of these mutants on MCP assembly. Notably, only strains containing PduA mutants capable of forming hydrogen bonds (R79R and R79K) are also capable of forming MCPs (Fig. 6C). These strains had significantly more puncta than all other mutant strains (p < 0.001) (Fig. 6C). This includes strains that were demonstrated to have lower, but detectable, levels of assembly (R79L, R79F, R79Y, R79W) (Fig. 3E). This is likely due to the fact that these strains are not able to form hydrogen bonds with the adjacent hexamer and therefore have lower binding stabilities. To confirm these results, we attempted to purify MCPs from a number of the PduA point mutant strains which did not show MCP assembly in the GFP encapsulation assay. However, we were unable to purify MCPs from these strains, as evidenced by the lack of the standard MCP banding pattern by sodium dodecyl sulfate gel electrophoresis (SDS-PAGE) (SI Fig. S13). Together, these results demonstrate that only PduA variants capable of forming hydrogen bonds enable MCP assembly. This validates the results shown in Fig. 4A which indicates that reduction of ε_hh_ disrupts MCP assembly and leads to no assembly.

## Discussion and Conclusion

Combined multi-scale MD simulations and *in vivo* mutation experiments show that hydrogen bonding and Coulomb interactions of arginine residues on the hexagon edges (ARG79) of PduA play a crucial role in native PduA self-assembly. In our experiments, mutations of ARG79 to most other residues negatively affect the assembly of Pdu proteins into tubes, cones or polyhedral compartments. From the fourteen mutations tested by All Atom (AA) MD simulations we found that those that have smaller bending and twisting angles are shown experimentally to be more likely to assemble into extended structures.

We determine the shape of these extended structures (tubes or cones) and confirm that small twisting and bending angles promote their formation by performing coarse-grained (CG) MD simulations. The CG model also determines the conditions to form polyhedral and icosahedral shells. Bacterial microcompartments (BMC) contain hexameric (BMC-H), pseudohexameric trimers (BMC-T), and pentameric (BMC-P) structures, which are considered in our CG model. Our CG model includes twisting and bending by considering the shape, the position of the charges and specific short-range interaction (including hydrogen bonding and hydrophobic interaction) of Pdu proteins in the form of BMC-H, BMC-T and BMC-P structures. In agreement with our experimentally observed trends, the CG model predicts that small bending and twisting angles facilitate PduA self-assembly into extended cylinders. However, the agreement is only qualitative because the CG bending angle used in our CG simulations is overestimated; that is, tends to be larger than the equilibrium bending angle obtained in the AA simulations, which include more detailed interactions.

According to our CG MD simulations, BMC-H proteins that maintain proper bending and twisting angles can form MCPs when BMC-P proteins and BMC-T proteins are present and BMC-P interactions with BMC-H are strong enough. For BMC-P/BMC-H mixtures that can form MCPs, the number ratio of BMC-H to BMC-P determines the size and asphericity of assembled shells, with a higher ratio of BMC-H leading to larger, more elongated shells. We found that increasing the content of BMC-T increases the asphericity of assembled structures (SI Fig. S6) because BMC-T prefers smaller bending angles. In some assemblies, pentameric units are stuck in 6-coordinated sites, which is against the expectations of elasticity theory, since pentameric dislocations should repel one another ^12^. This is likely due to local kinetic traps that are not overcome by annealing (see methods).

Other morphologies including “samosas” and cylinders arise in our CG simulations and theoretical arguments by modifying the hexamer and pentamer interactions via mutations. These morphologies are robust in simulations using other CG bending angles (see SI Figs. S4 – S5) and are also observed using a thermodynamics model that includes interactions, bending energies and concentrations of the components via chemical potentials. Our experiments demonstrate that reducing the hexamer-hexamer interaction strength abolishes MCP assembly in GFP encapsulation assays, as predicted by our CG MD simulations and theory. Our work explains the necessary conditions for BMC domain proteins to assemble into closed MCPs, as well as cylinders, and suggests that it is possible to systematically modify the shape of the resulting assembled structures.

## Materials and Methods

### All atom (AA) molecular dynamics (MD) simulations

The PduA hexamers (PDB id: 3ngk) in the AA MD simulations are downloaded from the PDB and solvated in water containing 100 mM NaCl and 4 mM MgCl_2_. Afterwards, the system undergoes a short constant volume, temperature (NVT) equilibration of 100 ps with the backbone restrained. Then, restraints are released for one PduA hexamer, while the other still has backbone restraint, and constant pressure, temperature (NPT) production simulations of 100 ns and umbrella sampling simulations are performed. For native PduA, an NPT extended simulation of 200 ns is performed, the PduA-PduA angle calculated are similar to 100 ns simulations. More details for AA MD simulations are in the SI.

### Coarse-grained (CG) MD simulations

The BMC-H, BMC-T, and BMC-P models include short-range attractive interactions modeled by a Lennard-Jones-Gauss potential on a pair of attractive sites on the side of CG proteins (Fig. 2A), given by

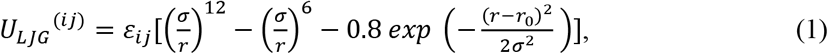

where σ = 1.0 nm is the bead size, *r*_0_ is 1.2 nm roughly corresponding to the length of arginine side chain and *i j* refers to the three species, p for pentamer (BMC-P), h for hexamer (BMC-H) and t for trimer (BMC-T), *ε*_*ij*_ is the binding energy between species *i* and *j*, where we choose the hexamer-hexamer interaction based on the AAMD simulation (*ε*_hh_ = *ε*_hh,AA_ + *μ*, *μ* is the CG chemical potential). To reduce the enormous parameter space, the value of *ε*_*ij*_ is set equal for the BMC-H and BMC-T proteins. The consequence of this assumption is discussed in the SI. *ε*_*ph*_ is unknown and left as a free parameter (see SI Table S2). In addition to the hydrogen bonding, the proteins’ charges (Fig. 2A right column) interact via a screened Coulomb potential with Debye length of 0.91 nm. The resulting total hexamer-hexamer interaction from two sites is then about −7k_B_T (or - 4.4 kCal/mol) per edge. The spherical cargo is a generic model for enzymes encapsulated in MCPs, they are attracted to BMC-H and BMC-T proteins at six sites by a standard 12-6 Lennard-Jones (LJ) potential with *ε* = 1.8, resulting in a total cargo/shell protein binding energy of ~ −7.5 kBT. The energy scale for this LJ potential was selected such that the cargo can bind to BMC-H and BMC-T, but the MCP structures are not heavily deformed by the strength of the cargo/shell protein attraction. Note that the hexamer-hexamer interaction energy in the CG model is smaller than that calculated from AA simulations because the protein concentration in CG simulations is higher (See SI appendix for details of MD simulations).

### Plasmid and strain creation

All plasmids and strains used in this study are listed in SI Tables S3 and S4. For modifications to the Pdu operon in LT2, λ red recombineering was used as previously described (see SI “Extended Materials and Methods”).

### PduA self-assembly assay

The PduA self-assembly assay was carried out as previously described (see SI “Extended Materials and Methods”). Once cultures were grown to saturation, strains were imaged using phase contrast microscopy. All images were adjusted equally for brightness and contrast and were cropped to an area of 500 x 500 pixels. Measurements of cell length were done using the segmented line tool in ImageJ as described previously.^22^

### Western blot

Western blots were done on cell cultures expressing FLAG-tagged PduA variants (see SI “Extended Materials and Methods”).

### GFP encapsulation assay

The GFP encapsulation assay was carried out to measure differences in MCP assembly between PduA variant strains as described in previous manuscripts (See SI “Extended Materials and Methods”).

### Thermodynamic theory

In this section, we describe the thermodynamic model that explains the morphologies observed in CG MD simulations. Consider *N*_*s*_ protein monomers (corresponding to BMC-H hexamers in the MD simulations) and *N*_*c*_ enzyme cargo initially placed in the simulation box. When the system reaches equilibrium, the monomers are distributed either as free monomers (*N*_0_) or as subunits of an oligomer (*N*_*q*_) that contains q monomers. The total number of monomers is conserved by the relation *N*_*s*_ = *N*_0_ + *qN*_*q*_. Similarly, the free (*f*) and encapsidated (*N*_*ce*_) cargo is subject to the conservation constraint *N*_*c*_ = *N*_*cf*_ + *N*_*ce*_. The free energy thus can be written as a sum of ideal free energy of each component and the interaction energy arising from shell bending energy, monomer-monomer binding energy and the cargo-monomer interaction:

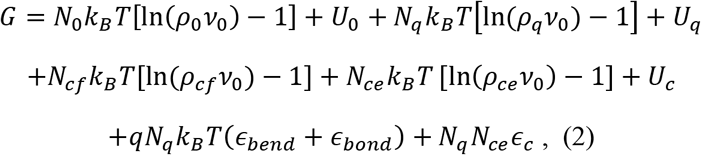

where *v*_0_ denotes the standard volumetric cell size (*a*^3^), with *a* approximated by the monomer size, and *∊*_*bend*_, *∊*_*bond*_ and *∊*_*c*_ represent the bending energy, binding energy per monomer and cargo-monomer interaction, respectively. Note that we have assumed the total internal energy for monomers (*U*_0_+*U*_*q*_) and cargo (*U*_*c*_) are constant.

Minimizing the free energy with respect to the number of free monomers we obtain the equilibrium condition:

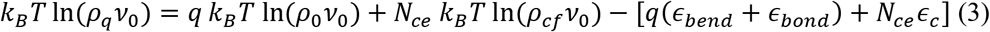

and the law of mass action for the particle density ^42^

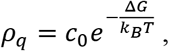

where Δ*G* = *q∊* − *qμ*_0_ − *N*_*ce*_*μ*_*c*_ is the formation energy with the excess energy given by 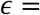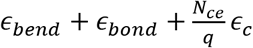 and the monomer and cargo chemical potentials are *μ*_0_ = *k*_B_*T* ln(*ρ*_0_*v*_0_) and *μ*_*c*_ = *k*_*B*_*T* ln(*ρ*_*cf*_*v*_0_), respectively.

To investigate the formation of the shell, we incorporate the line tension ^43, 44^so that

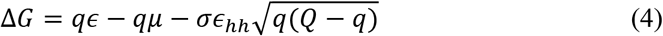

where 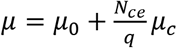 and 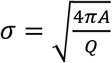, *Q* is the monomer number of a complete shell and 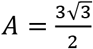 is the area factor of hexamer. Since the main contribution to the bending energy comes from the block geometry, it is reasonable to set the corresponding rigidity (*k*_*b*_) as a piece-wise function, i.e., *k*_*b*_ = 0 when no geometry overlaps and *k*_*b*_ is finite when overlap occurs. In such a case, only the corners of samosa shape and cylinder shape assemblies are subject to the bending penalty, with the local curvature defined by the cargo size (*r*_*c*_). Thus, we write the formation energy for each morphology as

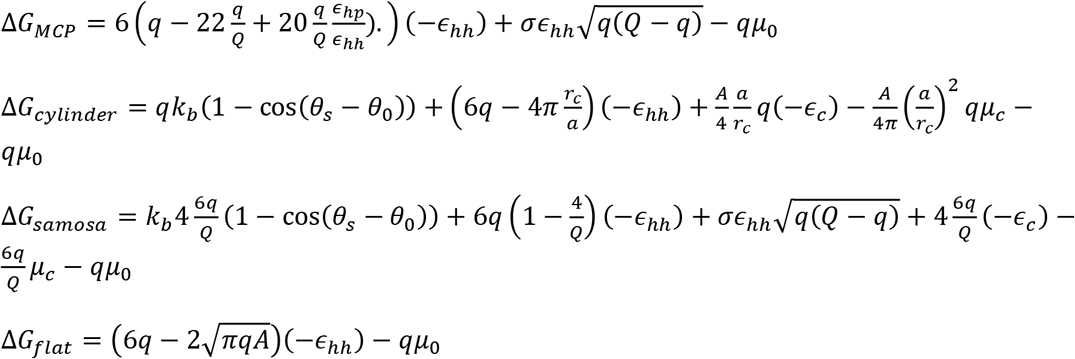

where *θ*_*s*_ is the angle between normal vectors of connected subunits and *θ*_0_ is the spontaneous angle determined by the intrinsic properties of the protein. (See SI Table S5 for the list of thermodynamic parameters).

## Acknowledgments

Y.L., S. L. and M.O.d.l.C. were supported by Department of Energy Award DE-FG02-08ER46539 and thank the Sherman Fairchild Foundation for computational support and Baofu Qiao for helpful discussions. D.T.E. and C.E.M. were supported by Army Research Office Award W911NF-19-1-0298. N.W.K. was supported by the National Science Foundation Graduate Research Fellowship Program (DGE-1842165), and by the National Institutes of Health Training Grant (T32GM008449) through Northwestern University’s Biotechnology Training Program.

