## Supplementary Appendix for "Computational and experimental approaches to controlling bacterial microcompartment assembly"

* Corresponding authors

**Table of contents:**

Extended Computational Methods

Extended Experimental Materials and Methods

Figure S1-13

Table S1-5

Legends for Supplementary video 1 and 2

References

**Supplementary Information Text**

**Extended Computational Methods**

**Atomistic molecular dynamics simulation methods.** All atom molecular dynamics simulations were performed using the GROMACS simulation package (version 2016.3)(1, 2),using Amber ff99SB-ILDN force field (3). A tetragonal box of 24*18*16 $nm^3$ was used for constant volume, constant temperature (NVT) simulations of 2 PduA hexamers, with periodic boundary conditions. The proteins were equilibrated for 100 ps in a constant volume, constant temperature (NVT) simulation with the backbones restrained with a force constant of 1000 kJ/mol/$nm^2$. In this work all simulations were performed at the temperature of 300 K unless otherwise noted. Then we released the restraints of one hexamer such that it was free to move or rotate to explore all possible configurations, while still ensuring that all atoms associated with protein molecules were at least 5 nm away from box edges. We then performed 100 ns constant pressure and temperature (NPT) simulations to obtain the angle between the norm of two hexamer planes. An additional 100 ns simulation NPT simulation was performed for the native PduA (pdb id: 3ngk). The angles plotted in Fig. 1 are defined by the norm of hexamer planes defined by the $\alpha$ carbon on 3 of the 6 arginines at $79^{th}$ position. To calculate the angle, the PduA hexamer whose backbone was restrained was taken as a reference frame and the unit vector of the other PduA was projected onto this cartesian frame to obtain bending and twisting angles. A summary of all the calculated angles is shown in Table S1. Mutations to the PduA structures were performed using the PyMol software.

To probe the PduA hexamer binding energy, steered simulations were employed to generate approximately 18 snapshots along the reaction coordinate (the center of mass distance between the two proteins) at an interval of about 1 Å. 10 ns constant pressure (NPT) umbrella sampling simulations were then performed for each snapshot using a spring constant of 2000 kJ/mol/$nm^2$. Weight histogram analysis was used to calculate the potential of mean force between the two proteins. The error is estimated by the bootstrapping tool implemented in g_wham (4), with 100 bootstraps. The potential of mean force is shown in Figure 1 A of main text. In the sampling of PduA hexamer binding, we took advantage of the fact that arginine 79 is always the binding location, and the natural docking orientation of the two proteins falls into a narrow angular range, allowing us to have decent sampling using 10 ns umbrella windows. The umbrella sampling simulations are repeated for native PduA and results are similar. For PduA R79 mutants, this specific short-range interaction is lost, and many different docking orientations thus become available. Therefore, the PMF for these mutants are not calculated.

**Correlation of BMC-H dimerization and hydrogen bonding.** To investigate the nature of the interaction between BMC-H proteins, we used the H-bond calculator in VMD (5) , with a cutoff distance of 3.3 Å and cutoff angle of 30°, to count the number of hydrogen bonds in umbrella sampling simulations. The number of H-bonds correlates well with the attachment state of the two PduA hexamers(Supplementary video S2). We also calculated the H-bonds on the $79^{th}$ residue for hydrophilic residues, summarized in Table 1 in the main text. Only the native PduA and the PduA R79K mutant are able to form hydrogen bonds between the $79^{th}$ residue with the main chain carbonyl oxygen of VAL 25. The lysine at position 26 in PduA (LYS26) has also been shown to form H-bonds (6). However, in our simulations, we find that this residue only forms H-bonds when the H-bond cutoff is increased to 3.5 Å, and even then these H-bonds only have an occupancy of 11% .

**Coarse-grained model.** A graphical illustration of the CG model is shown in Fig.2 in the main text. The bottom sides of the 3 proteins multimers (hexamer, trimer, pentamer) are smaller than the top sides of these multimers (see all-atom PDB structures), forming an inclination angle. To account for this in our coarse-grained model, the side of each multimer is inclined at an angle $\theta_i$, which is related to the 2-plane angle $\theta$ in the main text by $\theta = 2\theta_i$. Charge beads are placed on the CG structures based on the total charge and dipole moment of the protein. PDB ids used for coarse-graining are 3ngk for BMC-H, 4fay for BMC-T and 4i7a for BMC-P. Cargo spheres are intended to represent a generic enzyme, and have been constructed to have a similar diameter and charge to the diol dehydratase-cyanocobalamin complex, PDB id 1egm, which is meant to serve as an analog to the diol dehydratase natively encapsulated in Pdu MCPs. Charge beads interact by a DLVO potential:

$$\beta U(r) = Z^2 \lambda_B \left(\frac{e^{\kappa a}}{1+\kappa a}\right)^2 e^{-\kappa r}/r \qquad \text{(Eq. S1)}$$

The protein charges are calculated using PROPKA (7, 8) The total net charges at pH=7 are 3ngk: 0 e, 4fay: -10 e, 4i7a: -10 e. BMC-H proteins have a net charge close to 0 at pH 7, but the top and bottom side of the are charged differently at this pH (Fig. S2). This charge distribution is represented by positive and negative beads shifted in the vertical direction. The hexamers, pentamers, trimers, and cargo are treated as rigid bodies (9). It is important to note that both shape asymmetry and charge distribution are needed for the building blocks to form closed compartments. If the charges are removed, the proteins form large, slightly curved sheets. The ARG79 interaction sites, which is critical for dimerization, are modeled by the Lennard-Jones-Gauss (LJG) potential

$$U_{LJG}{}^{(ij)} = \varepsilon_{ij}\{\left(\frac{\sigma}{r}\right)^{12} - \left(\frac{\sigma}{r}\right)^{6} - \alpha \exp\left(-\frac{(r-r_0)^2}{2\sigma^2}\right)\}. \qquad \text{(Eq. S2)}$$

The Gaussian term in this potential serves as a modifier to the slope of the potential, facilitating the search for binding sites. $\alpha$ takes the value of 0.8 such that the profile of LJG potential resembles the potential measured by atomistic umbrella sampling. Constant volume CG simulations with periodic boundary conditions are performed using the Langevin integrator implemented in HOOMD (10, 11).The simulations are equilibrated at 300 K (Corresponding to T=1 in LJ temperature) for $1 \times 10^7$ time steps and annealed gradually to 345 K in a cycle. The simulation box is constructed by replicating a repeating unit box 5 times in x, y and z direction. The simulation box for the simulation in Figure 2 is 26.25*26.25*15.25 $nm^3$, we note that this result in a concentration much higher than *in vivo* or lab conditions to reach assembly faster. All assembly shapes are obtained after 3 cycles. There is no significant difference between the assembled shape in cycle 2 and 3, so we have good confidence that we have obtained the equilibrium structures.

To alter the twisting angle between BMC-H structures in the CG model, we shift the attractive sites representing the ARG79 hydrogen bond in the plane normal direction by

$$\Delta h = \frac{1}{2} l_s \tan\theta_t, \qquad \text{(Eq. S3)}$$

where $l_s$ is the distance between the arginine site (pink bead in figure2 of main text) and the binding sites (purple beads) on one PduA hexamer.

**Parameterizing the coarse-grained model of BMC-T proteins.** The PduB trimer (PDB id 4fay) has an almost perfect hexagonal symmetry, which is the same symmetry as PduA hexamers. Thus, it is difficult to intuitively predict the role of PduB in MCP assembly. In our CG model, the BMC-T is differentiated from the BMC-H by an extra layer of beads on its top side, representing the fact that these BMC-T trimers are typically thicker than BMC-H hexamers. The BMC-T structures therefore have slightly different bending angle and bending rigidity, which affects the overall shape of MCPs. This is predicted by previous theoretical work (12), as well as experimental studies that have found that the mean curvature of tubes formed by PduA and PduB is larger than that of tubes formed by PduA alone (13). We study the effect of BMC-T presence on MCP formation by changing the number ratio of BMC-T to BMC-H and the interaction strength of BMC-T with other shell proteins. At a constant total number of BMC-H and BMC-T proteins, we increase the number of BMC-T proteins in a unit cell from 0 to 4 to 6 (Figure S6 A-C). With no BMC-T, the system forms more symmetric MCPs. In the case where $N_{BMC-H}$:$N_{BMC-P}$=5:3, they form T=3 icosahedral shells (see Figure 2B in main text). When number of BMC-H proteins equal to that of BMC-T proteins, the MCPs are more aspherical and elongated. At $N_{BMC-H}$:$N_{BMC-T}$=2:6, assembled shells are more irregular. Having stronger attractive interactions between BMC-T proteins than BMC-H proteins ($\varepsilon_{tt} = 1.2\varepsilon_{hh}$) causes BMC-T proteins to preferentially assemble into quasi-closed polyhedral structures (Figure S6D). Since in native Pdu MCPs the shells are closed structures, BMC-T should not have a much stronger attraction than BMC-H, which is in line with our assumption.

**Effect of stoichiometry on assembled shapes.** Given the vast parameter space of this multi-component system, we focus here on the ratio of BMC-H proteins to others, for which we have more experimental data for comparison. In the simulation box, we have 1 BMC-P, 2 BMC-T, 2 spherical enzymes in one repeating unit, and vary the number of BMC-Hs, $\mathrm{n_{BMC-H}}$. The unit cell is replicated 125 times so the total number of BMC-H $\mathrm{N_{BMC-H}} = 125 \times n_{\mathrm{BMC-H}}$. In the range of 4 to 8 BMC-Hs, there is a small trend of increasing BMC size and the corresponding asphericity (14) (Figure S7). This is consistent with the fact that when $N_{\mathrm{BMC-H}}$ approaches infinity, cylinders are observed both in experiments and simulations. In the simulations with $N_{\mathrm{BMC-H}} = 16$ and 32, the assembled size appears to be bigger, but until the end of $3 \times 10^7$ time steps of simulations, the proteins cannot form closed compartments.

**Effect of $\boldsymbol{\theta_i}$ on assembled shapes**. We show that the CG bending angle is approximately $2\theta_i$ by running a CG MD simulation of two BMC-H proteins of different inclination angle $\theta_i$ and keeping track of the angle formed by the norm of the two hexagons. The distribution is fitted to a gaussian function and the mean angle is plotted against $2\theta_i$, as shown in Figure S4A. We also investigated the effect of CG bending angle on the geometry of self-assembled structures. We found that BMC proteins with bending angles of $\theta_i = 15$ and with $\theta_i = 5$ form similar morphologies to those observed in simulations using $\theta_i = 25°$. We determined the phase diagram for $\theta_i = 15°$ analogously to Figure 4 in the main text. The two-phase diagrams demonstrate similar behavior with MCPs formed in the upper middle region ($3.4 < \varepsilon_{hh} < 4.2$). The main difference is that for $\theta_i = 15°$, cylinders have larger radius. In the lower right region where cylinders are expected, another rod like structure with defects is observed (Figure S2D). Also, cones with zero gaussian curvature like cylinders are observed (Figure S4E).

### Extended Experimental Materials and Methods

**Plasmid and strain creation.** All plasmids and strains used in this study are listed in Supplemental Table 2-3. Plasmids were created using Golden Gate cloning. Briefly, a modified pBAD33t expression vector was created which includes the first 74 residues of PduA, followed by a GFP insertion cassette, flanked by BsaI cut sites. DNA oligos containing PduA point mutations were ordered from Sigma Aldrich, PCR amplified with primers containing BsaI cut sites, and then cloned into the pBAD expression vector using established methods (16). Plasmids were sequence verified using Sanger sequencing from Quintara.

For modifications to the Pdu operon in LT2, λ red recombineering was used as previously described (17). Briefly, the cat-sacB selection cassette was PCR amplified to contain homology to a specific Pdu locus and integrated. *PduA* variants were PCR amplified from the previously described expression vector constructs to contain homology to the *pduA* locus. These homology-containing PCR products were electroporated into

LT2 strains containing the integrated cat-sacB cassette. Induction of the λ red recombination system allowed for *pduA* variants to integrate at the *pduA* locus in place of the cat-sacB cassette. Strains were plated on 6% sucrose plates to test for sucrose sensitivity, and positive strains were sequence verified using Sanger sequencing from Quintara.

**PduA self-assembly assay.** The PduA self-assembly assay was carried out as previously described (18). Briefly, PduA variants were cloned into modified pBAD33t expression vectors as described above. Strains were stored as 15% glycerol stocks at -80°C until use and were restreaked onto Lysogeny broth – Miller (LBM) plates containing 34 µg/mL chloramphenicol (cm). Restreaks were grown for 16 hours at 37°C, and colonies were used to inoculate 5 mL cultures of LBM + cm in 24-well blocks. Inoculated cultures were grown overnight (16 hours) at 37°C, 225 RPM. After overnight growth, cultures were used to inoculate fresh 5 mL LBM + cm cultures 1:100. These cultures were grown for 90 minutes (or to an $OD_{600}$ of 0.2-0.5) at 37°C 225 RPM and then induced by adding 50 µL of 20% (w/v) arabinose to a final concentration of 0.2%. Induced cultures were then incubated at 37°C, 225 RPM for 4-6 hours or until cultures were saturated.

Once cultures were grown to saturation, strains were imaged using phase contrast microscopy. For each strain, 1.48 µL of culture was prepared on Fisherbrand™ frosted microscope slides and 22 x 22 mm, #1.5 thickness cover slips. Imaging was performed with a Nikon Eclipse Ni-U upright microscope containing a 100X oil immersion objective lens. Images were collected using the Andor Clara digital camera and were initially processed using NIS Elements software (Nikon). Further image analysis was done using ImageJ. All images were adjusted equally for brightness and contrast and were cropped to an area of 500 x 500 pixels. Measurements of cell length were done using the segmented line tool in ImageJ as described previously.

**SDS-PAGE and western blot.** Western blots were done on cell cultures expressing FLAG-tagged PduA variants. For each variant, 1 mL of induced culture was pelleted at 5k x G, the supernatant was removed, and the pellets were stored at -20°C until use. Pellets were then resuspended to an $OD_{600}$ of 3.0 in lysis buffer containing sodium dodecyl sulfate (SDS) (25 mM Tris base, 192 mM glycine, and 1.1% SDS). Samples were combined 3:1 with 4X concentrated Laemmli buffer at heated for 5 minutes at 95°C. These boiled and denatured samples were then loaded onto a 15% Tris-glycine SDS-PAGE gel and run at 130V for 80 minutes in preparation for western blotting, or at 120V for 90 minutes for Coomassie Brilliant Blue R-250 staining. The gel was then prepared for wet transfer and transferred onto a PVDF membrane at 90V for 11 minutes. After transfer, membranes containing protein were blocked for 1 hour at room temperature with gentle rocking in TBS-T (20 mM Tris (pH 7.5), 150 mM sodium chloride, 0.05% (v/v) Tween-20) and 5% (w/v) dry milk. Membranes were incubated overnight (~16 hours) at 4°C with gentle rocking in TBS-T + 1% (w/v) dry milk with 1:6666 mouse anti-FLAG primary antibody. These membranes were then washed 4x with TBS-T. Washed membranes were incubated with 1:1000 goat anti-mouse antibody conjugated with horse radish peroxidase (HRP) for 90-120 minutes at room temperature with gentle rocking. The membranes were washed again 4x in TBS-T before development with SuperSignalTM West Pico PLUS Chemiluminescent Substrate (Thermo Fisher Scientific). Images were collected and initially processed on the Bio-Rad ChemiDoc XRS+ System. Densitometry was done using Image Lab software for relative PduA expression measurements.

**GFP encapsulation assay.** The GFP encapsulation assay was carried out to measure differences in MCP assembly between PduA variant strains as described in previous manuscripts (18). Briefly, overnight cultures of modified LT2 strains containing pBAD33t-ssD-GFPmut2 plasmid (CMJ069) were prepared as described above in 5 mL LBM cultures in 24-well blocks. Cultures were grown overnight for ~16 hours at 37°C, 225 RPM. The overnight cultures were then subcultured 1:500 (10 µL) into 5 mL of LB-M with 0.02%

(w/v, final concentration) L-(+)-arabinose, 34 ug/mL cm, and 0.4% (v/v, final concentration) 1,2-PD to simultaneously induce GFP cargo and MCPs. Cultures were grown at 37 °C, (225 RPM) for a minimum of 6 hours post-subculture and then imaged with a Nikon Eclipse Ni-U upright microscope and C-FL Endow GFP HYQ bandpass filter for fluorescence images. An 80 ms exposure was used for all images, which were collected processed as described above. Images were adjusted using ImageJ software and cropped to a final area of 250 x 250 pixels.

**MCP purification.** MCPs were expressed by first restreaking strains from 15% glycerol stocks stored at -80°C on to LBM plates. Plates were incubated overnight (15-16 hours) at 37°C. Single colonies were picked from these plates and used to inoculate 5 mL cultures of LBM, which were grown for 12 hours at 37°C, 225 RPM. Cultures were then used to inoculate 200 mL No-Carbon E (NCE) (29 mM potassium phosphate monobasic, 34 mM potassium phosphate dibasic, 17 mM sodium ammonium hydrogen phosphate) cultures supplemented with 1 mM magnesium sulfate, 50 µM ferric citrate, and 55 mM 1,2-PD. Cultures were supplemented 1:1000 (200 µL into 200 mL of media) and were grown at 37°C and 225 RPM to an $OD_{600}$ of 1.5. MCPs were then purified as previously described using differential centrifugation (19). Briefly, cells are pelleted at 5k RPM for 5 minutes at 4°C, resuspended in a chemical lysis buffer (32 mM Tris, 200 mM potassium chloride, 5 mM magnesium chloride,1.2% (v/v) 1,2-PD, 0.6% (w/v) octylthioglucoside (OTG), 2.2 mM β-mercaptoethanol, 0.8 mg/mL lysozyme, and 0.04U/mL DNaseI), and incubated at room temperature for 30 minutes. Cell lysate was then clarified via centrifugation at 12k x *g*, 4°C, for 5 minutes. MCPs were then pelleted at 21k x *g* for 20 minutes. Pelleted microcompartments were resuspended at stored at 4°C in a buffered solution containing 50 mM Tris (pH 8.0), 50 mM potassium chloride, 5 mM magnesium chloride, 1% (v/v) 1,2-PD.

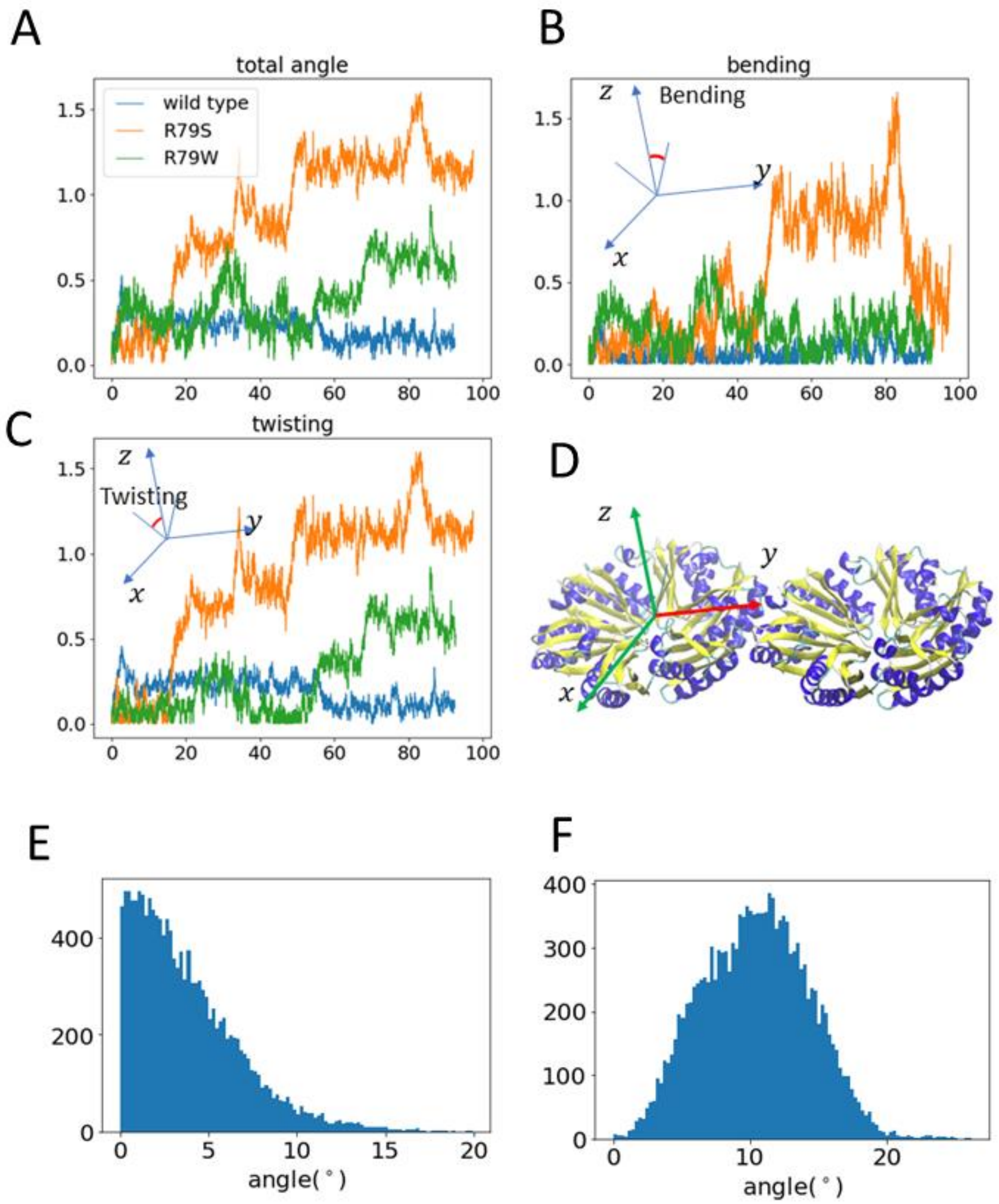


**Figure S1.** Orientation analysis of two PduA proteins in all atom simulations, comparing 3 characteristic types of protein: the native PduA has low overall angles and the highest stability; 2 R79W mutation hexamers have intermediate stability and have slightly larger twisting angles; R79S mutation is least stable and do not self-assemble according to our experiments. (A) The total angles of native PduA and two mutants, R79S and R79W. A Cartesian coordinate frame is established on the fixed BMC-H protein as shown in (D), and the surface normal of the other BMC-H is projected onto yz and xz plane to obtain the bending and twisting angles, respectively. The distribution of native PduA bending angle (E) and twisting angles (F).

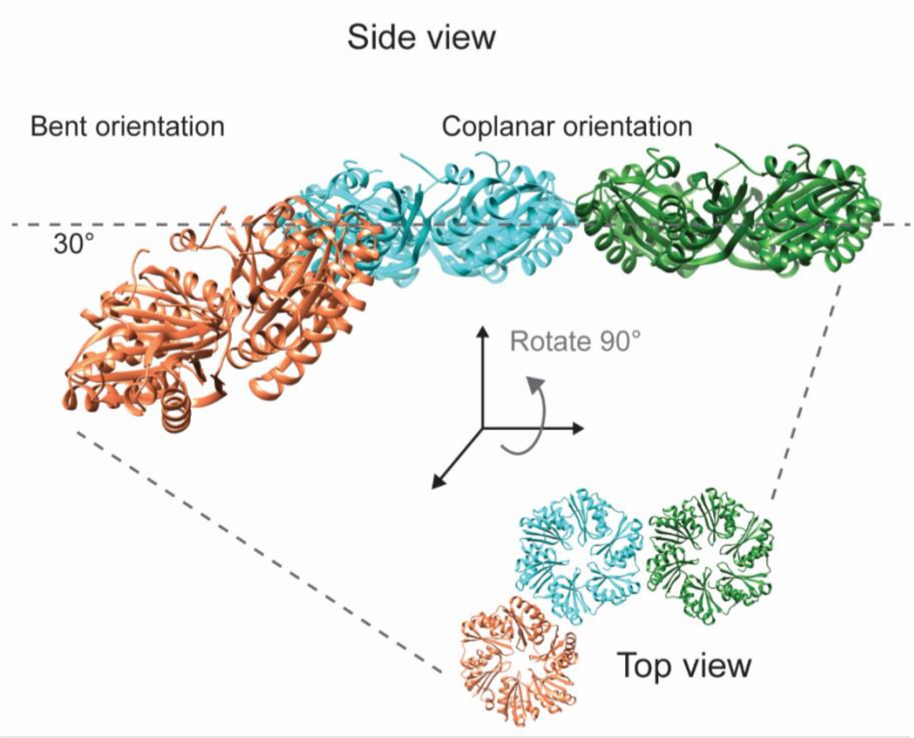


**Figure S2.** Bent and coplanar PduA interaction. PduA monomers (PDB ID: 3NGK) aligned with HO BMC monomers (PDB ID: 5V74) in both the bent (orange and teal hexamers) and coplanar (green and teal hexamers) orientations.

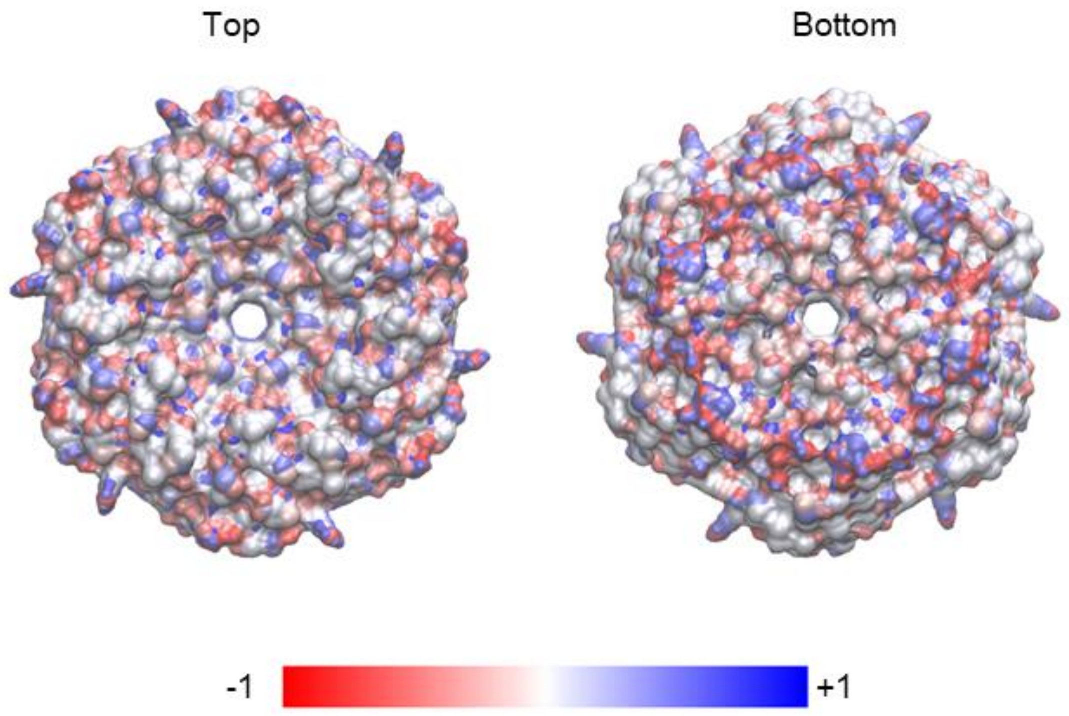


Figure S3. The charge distribution of PduA (pdb id: 3ngk) calculated by PDB2PQR server (7, 8), visualized using VMD (5).

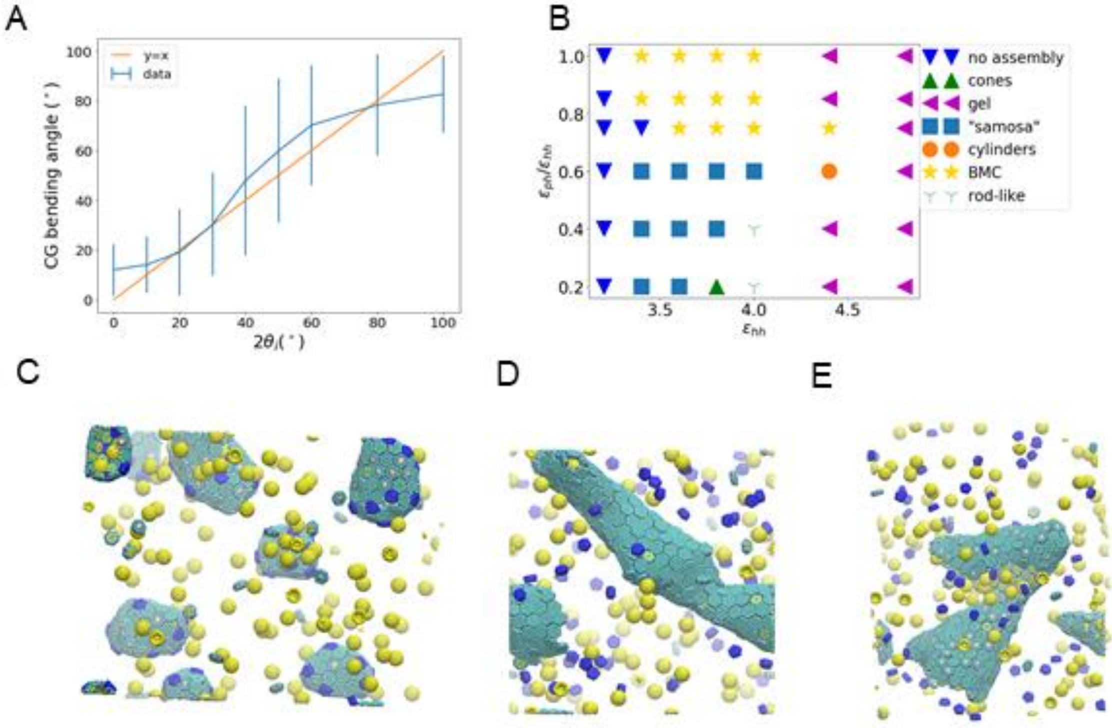


**Figure S4.** Analysis of coarse-grained (CG) bending angle. (A) CG MD simulations using different $\theta_i$ shows that the CG bending angle is ~$2\theta_i$. The number ratios are BMC-H : BMC-T : BMC-P : cargo=4:2:1:2 (B) Phase diagram calculated using $\theta_i = 15°$, corresponding to a cg bending angle of 30. (C), (D) and (D) are snapshots of CG simulations where the hexamers, pentamers and cargo are in green, blue and yellow, respectively: (C) MCP observed at $\varepsilon_{hh} = 3.8, \varepsilon_{ph} = 3.8$, (D) Rod-like structure observed at $\varepsilon_{hh} = 4.0, \varepsilon_{ph} = 0.8$, and (E) Cones observed at $\varepsilon_{hh} = 3.8, \varepsilon_{ph} = 0.76$.

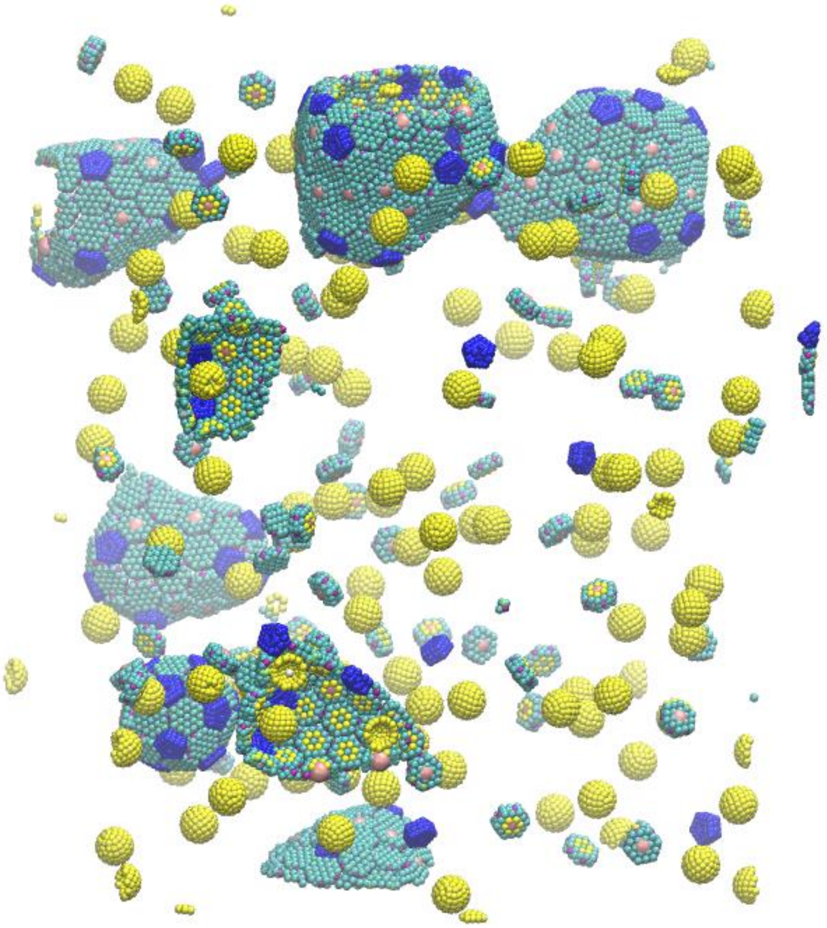

**Figure S5.** Snapshots of MCPs observed in CG MD simulations using the inclination angle $\theta_i = 5°$, CG bending angle of 10 (hexamers, pentamers and cargo in green, blue and yellow, respectively). The MCPs are qualitatively similar across different $\theta_i$ , suggesting that the shape of MCPs formed does not depend sensitively on $\theta_i$, but is primarily determined by stoichiometric ratios.

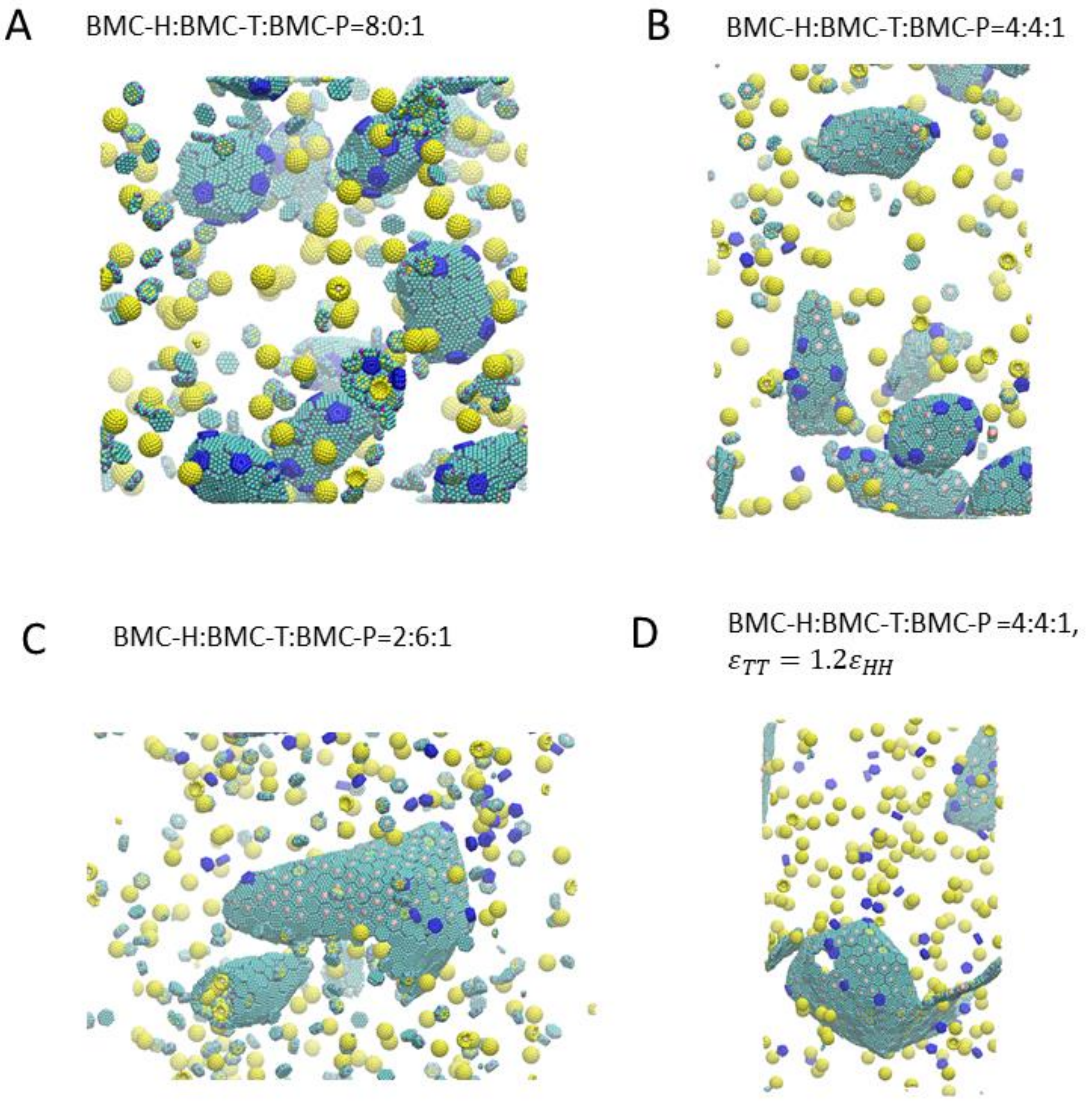


**Figure S6.** The role of BMC-T proteins in determining MCP shape. (A) to (D) are snapshots of CG MD simulations where the hexamers, pentamers and cargo are in green, blue and yellow, respectively. (A)~(C) An increasing number of BMC-T proteins relative to BMC-H proteins causes the assembly to shift from higher symmetry shells in (A) to more aspherical polyhedral shells in (C). The calculated asphericity are: (A) $0.15 \pm 0.09$, (B) $0.2 \pm 0.2$ and (C) $0.2$ (uncertainty nor available because there are two few data points). (D) Increasing the BMC-T/BMC-T interaction relative to the BMC-H/BMC-H interaction by a factor 1.2 causes BMC-T binding to be favored, having a similar effect as increasing the relative number of BMC-T proteins.

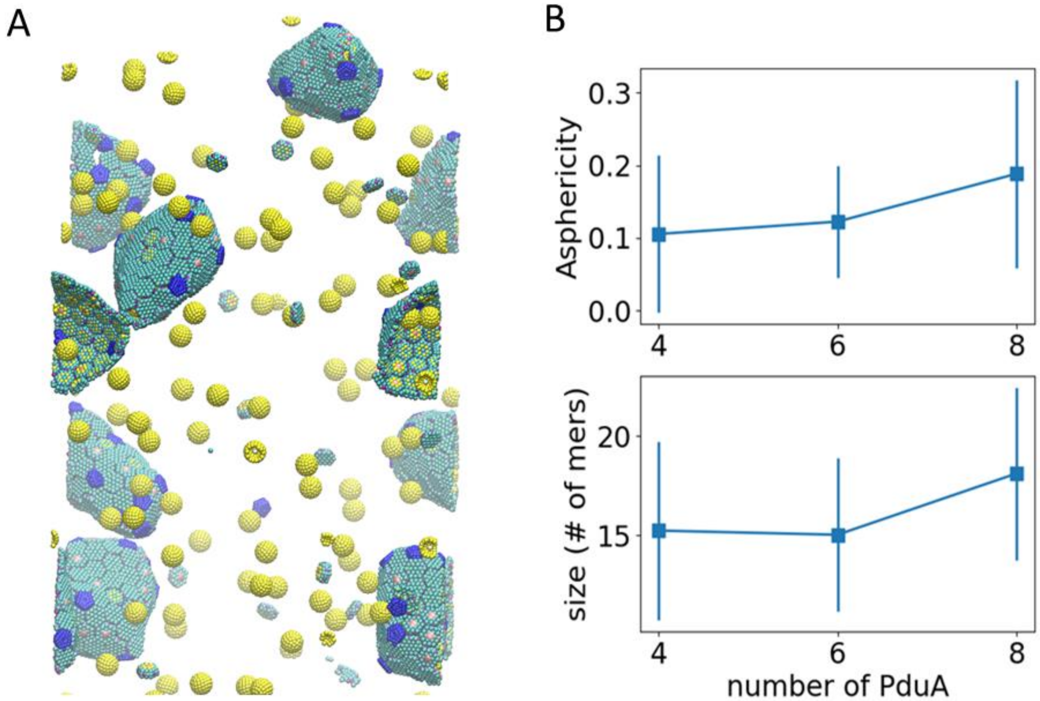


**Figure S7.** The change of MCP shapes for different BMC-H ratios. (A) The final CG simulation snapshot with $\mathrm{BMC-H:BMC-T:BMC-P = 8:2:1}$ (hexamers, pentamers and cargo in green, blue and yellow, respectively). (B) Asphericity and radius of gyration for varying $\mathrm{n_{BMC-H}}$ in a repeating cell. This cell is replicated 5 times in x, y and z direction. The error bar represents standard deviation of all observed MCPs in the equilibrium configuration.

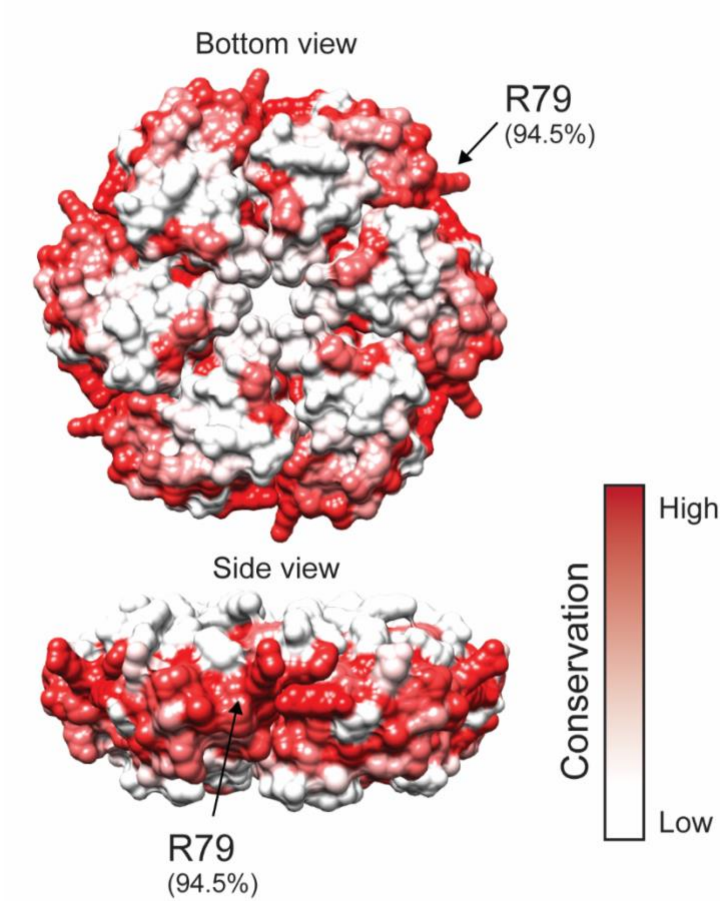


**Figure S8.** PduA conservation scores as determined by a multiple sequence alignment of 192 PduA homologs. Percent conservation scores overlaid is on the PduA structure, with ARG79. Red = high percent conservation, white = low percent conservation.

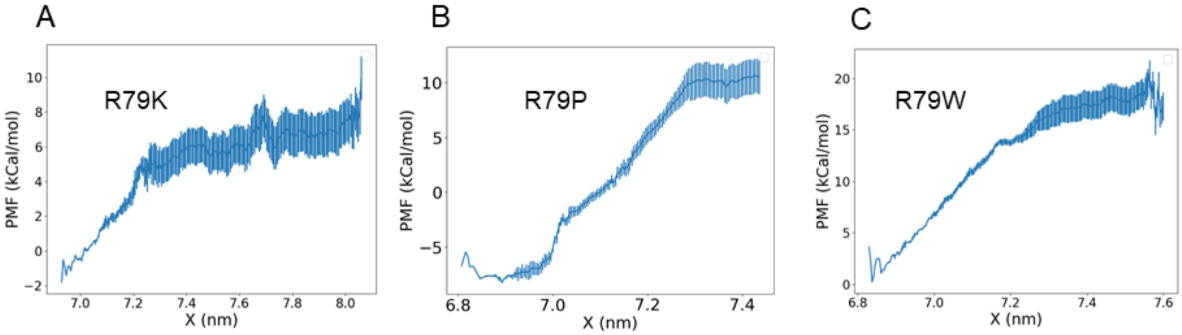


**Figure S9.** PMF of 3 selected mutations, R79K, R79P and R79W. R79K forms a hydrogen bond with the main chain VAL25 on the other hexamer and has similar binding energy to WT PduA. R79P and R79W both interact with neighboring PduA via hydrophobic interactions, but the hexamer-hexamer bending and twisting angles of R79P are much larger than R79W.

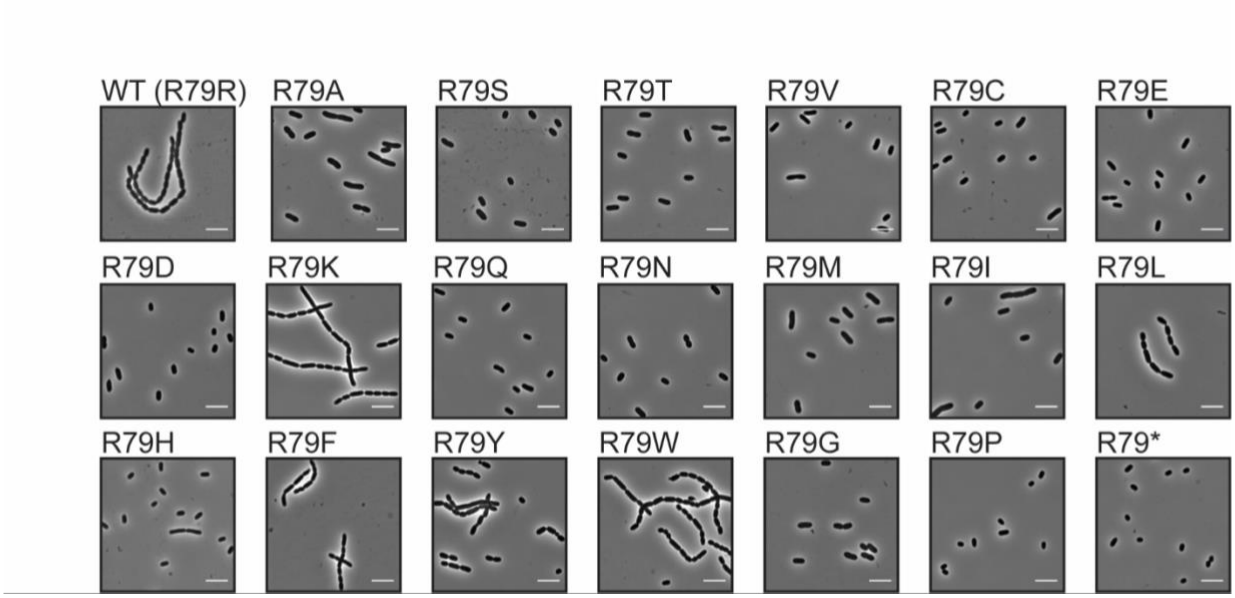


**Figure S10**. PduA self-assembly assay. Phase contrast microscopy images of cells overexpressing PduA variants (scale bar = 5 µm).

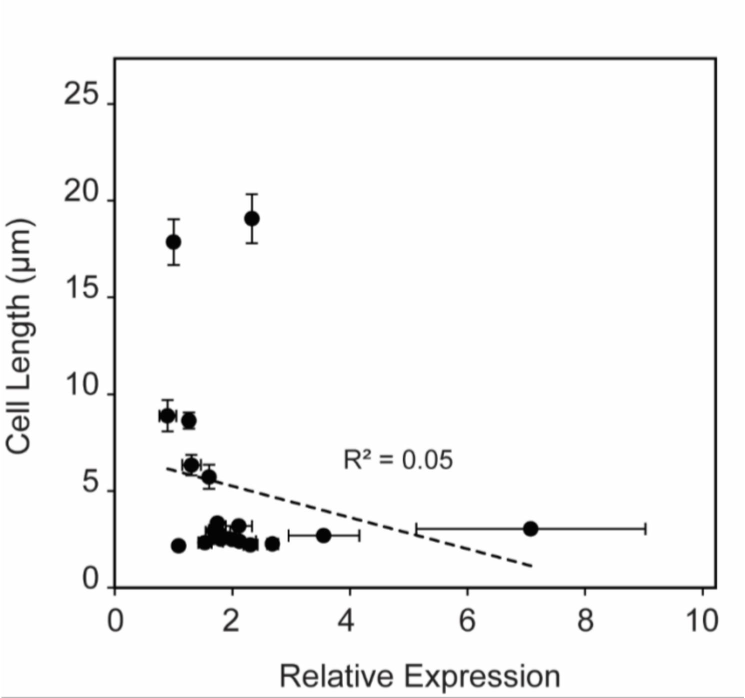


**Figure S11.** Correlation between cell length and protein expression. Cell length in μm (see Fig. 3E and SI Fig. S10) is compared to relative expression of PduA variants, as determined by semiquantitative western blot densitometry analysis (n = 3). Note that the correlation is weak ($R^2 = 0.05$).

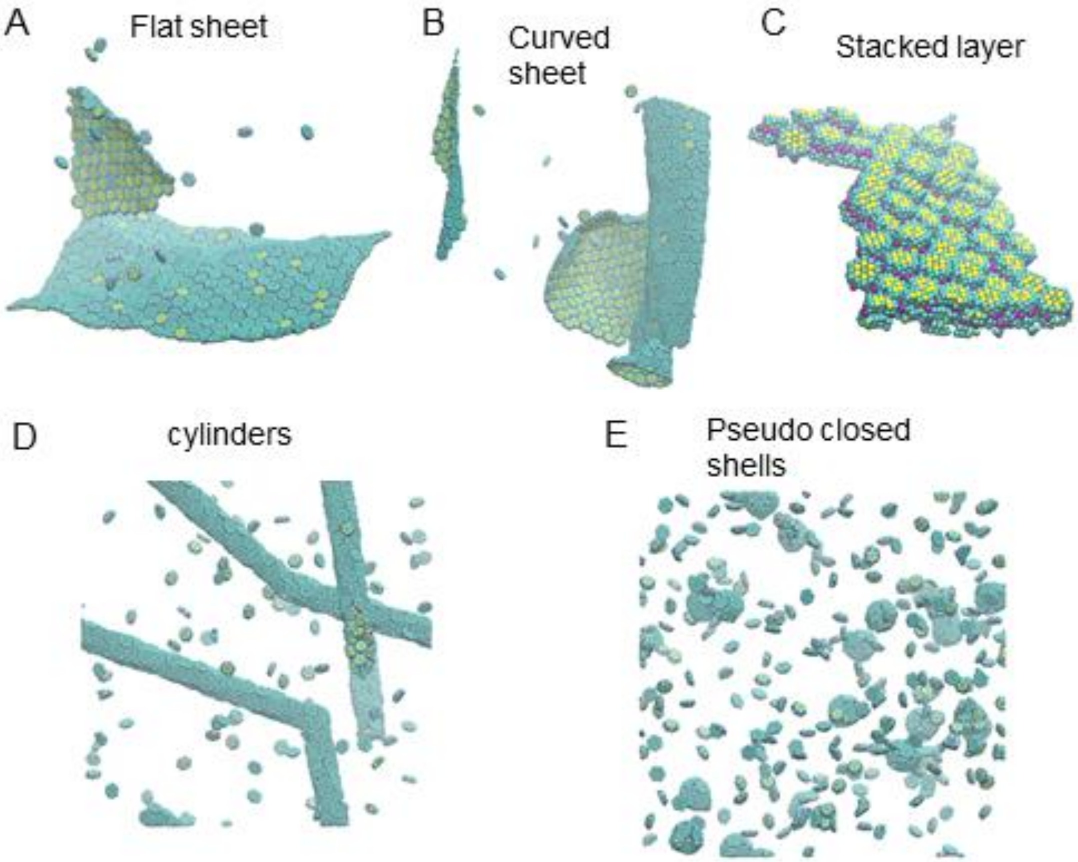


**Figure S12.** Examples of the morphologies discovered in the phase diagram of main text Fig. 3D. (A) Flat sheet observed at $\theta_b = 0, \theta_t = 0$. (B) Curved sheet observed at $\theta_i = 5, \theta_t = 0$. (C) Stacked layer aggregate at $\theta_i = 0, \theta_t = 80$. (D) Cylinders at $\theta_i = 25, \theta_t = 10$. (E) Pseudo closed shells observed at $\theta_i = 40, \theta_t = 0$.

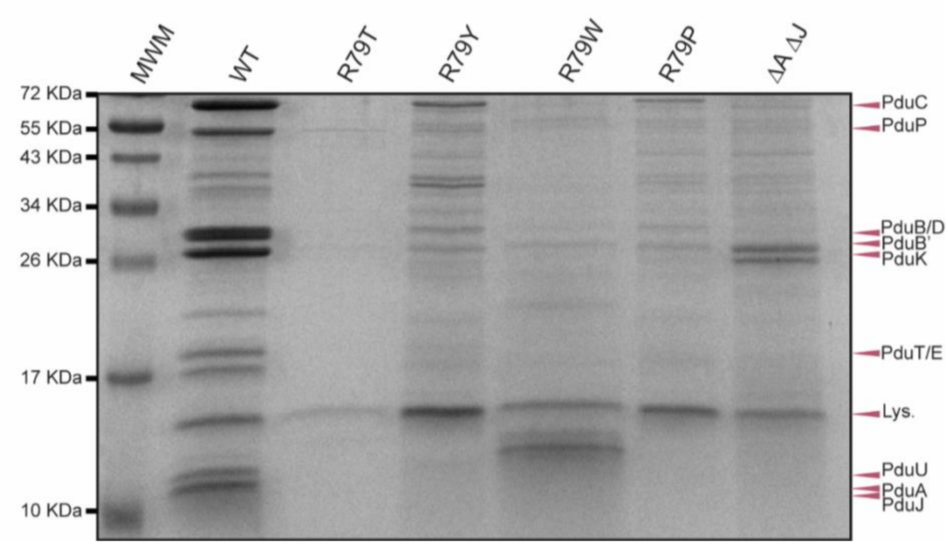


**Figure S13.** SDS-PAGE gel of MCPs. Presence of MCPs are determined by the standard MCP banding pattern (see WT lane). Note that the negative control ΔA ΔJ lane does not contain this standard banding pattern. MCP samples from strains with PduA variants also do not contain this banding pattern, indicative of malformed MCPs. "MWM" is the molecular weight marker. "Lys." is lysozyme used for cell lysis in the MCP purification method.

**Table S1**. Component and structural analysis of CG MCP using an initial ratio of BMC-H:T:P:cargo=4:2:1:2.

| MCP ID | BMC-H | BMC-T | BMC-P | total | Asphericity | Rg |
|---|---|---|---|---|---|---|
| 1 | 52 | 21 | 14 | 87 | 0.05 | 13.56 |
| 2 | 39 | 13 | 10 | 62 | 0.03 | 11.42 |
| 3 | 61 | 29 | 16 | 106 | 0.10 | 15.04 |
| 4 | 57 | 24 | 14 | 95 | 0.07 | 14.31 |
| 5 | 33 | 10 | 10 | 53 | 0.04 | 10.53 |
| 6 | 27 | 16 | 9 | 52 | 0.03 | 10.38 |
| 7 | 25 | 15 | 7 | 47 | 0.03 | 9.79 |
| 8 | 22 | 5 | 9 | 36 | 0.12 | 8.67 |
| 9 | 51 | 26 | 12 | 89 | 0.13 | 13.94 |
| mean | 40.78 | 17.67 | 11.22 | 69.67 | 0.07 | 11.96 |

**Table S2.** Detailed interaction parameters for simulations

| | $\varepsilon_{hh}$ | $\varepsilon_{ph}$ | $\theta_i$ | $\varepsilon_{pp}$ | $\varepsilon_{tt}$ | $\varepsilon_{th}$ |
|---|---|---|---|---|---|---|
| Fig. 2B | 4.0 | 3.4 | 25 | 0 | 4.0 | 4.0 |
| Fig. 2C-D | 4.0 | 3.4 | 25 | 0 | 4.0 | 4.0 |
| Fig. 2E | 4.0 | N/A | 25 | 0 | 4.0 | 4.0 |
| Fig. 4C | 3.8 | 1.9 | 25 | 0 | 3.8 | 3.8 |
| Fig. 4D | 3.8 | 2.28 | 25 | 0 | 3.8 | 3.8 |
| Fig S6 | 4.0 | 3.4 | 25 | 0 | 4.0 | 4.0 |
| Fig. S6D | 4.0 | 3.4 | 15 | 0 | 4.8 | 4.4 |

**Table S3.** Strains used for this manuscript.

| Strain | Organism | Genotype |
|---|---|---|
| DTE017 | *E. coli DH10b* | Wild type |
| DTE001 | *E. coli BL21* | Wild type |
| MFSS044 | *S. enterica serovar Typhimurium LT2* | Wild type |
| NWKs081 | *S. enterica serovar Typhimurium LT2* | *ΔpduJ* |
| NWKs083 | *S. enterica serovar Typhimurium LT2* | *ΔpduA ΔpduJ* |
| NWKs263 | *S. enterica serovar Typhimurium LT2* | *ΔpduA::pduA-R79A ΔpduJ* |
| NWKs264 | *S. enterica serovar Typhimurium LT2* | *ΔpduA::pduA-R79S ΔpduJ* |
| NWKs265 | *S. enterica serovar Typhimurium LT2* | *ΔpduA::pduA-R79V ΔpduJ* |
| NWKs266 | *S. enterica serovar Typhimurium LT2* | *ΔpduA::pduA-R79C ΔpduJ* |
| NWKs267 | *S. enterica serovar Typhimurium LT2* | *ΔpduA::pduA-R79E ΔpduJ* |
| NWKs268 | *S. enterica serovar Typhimurium LT2* | *ΔpduA::pduA-R79D ΔpduJ* |
| NWKs269 | *S. enterica serovar Typhimurium LT2* | *ΔpduA::pduA-R79K ΔpduJ* |
| NWKs270 | *S. enterica serovar Typhimurium LT2* | *ΔpduA::pduA-R79Q ΔpduJ* |
| NWKs271 | *S. enterica serovar Typhimurium LT2* | *ΔpduA::pduA-R79N ΔpduJ* |
| NWKs272 | *S. enterica serovar Typhimurium LT2* | *ΔpduA::pduA-R79M ΔpduJ* |
| NWKs273 | *S. enterica serovar Typhimurium LT2* | *ΔpduA::pduA-R79I ΔpduJ* |
| NWKs274 | *S. enterica serovar Typhimurium LT2* | *ΔpduA::pduA-R79L ΔpduJ* |
| NWKs275 | *S. enterica serovar Typhimurium LT2* | *ΔpduA::pduA-R79H ΔpduJ* |
| NWKs276 | *S. enterica serovar Typhimurium LT2* | *ΔpduA::pduA-R79F ΔpduJ* |
| NWKs277 | *S. enterica serovar Typhimurium LT2* | *ΔpduA::pduA-R79G ΔpduJ* |
| NWKs297 | *S. enterica serovar Typhimurium LT2* | *ΔpduA::pduA-R79T ΔpduJ* |
| NWKs298 | *S. enterica serovar Typhimurium LT2* | *ΔpduA::pduA-R79Y ΔpduJ* |
| NWKs299 | *S. enterica serovar Typhimurium LT2* | *ΔpduA::pduA-R79W ΔpduJ* |
| NWKs300 | *S. enterica serovar Typhimurium LT2* | *ΔpduA::pduA-R79P ΔpduJ* |

**Table S4.** Plasmids used for this manuscript.

| Name | Plasmid | Origin | Resistance |
|---|---|---|---|
| pSIM6(20) | λ Red system repressed by cI857 | pSC101 *repA*$^{ts}$ | Ampicillin |
| CMJ069 | pBAD33t-ssD-GFPmut2 | p15A | Chloramphenicol |
| CMJ138 | pBAD33t-PduA-FLAG | p15A | Chloramphenicol |
| NWKp004 | pBAD33t-PduA-entry vector | p15A | Chloramphenicol |
| NWKp015 | pBAD33t-PduA-R79A-FLAG | p15A | Chloramphenicol |
| NWKp019 | pBAD33t-PduA-R79D-FLAG | p15A | Chloramphenicol |
| NWKp023 | pBAD33t-PduA-R79S-FLAG | p15A | Chloramphenicol |
| NWKp024 | pBAD33t-PduA-R79T-FLAG | p15A | Chloramphenicol |
| NWKp025 | pBAD33t-PduA-R79V-FLAG | p15A | Chloramphenicol |
| NWKp026 | pBAD33t-PduA-R79C-FLAG | p15A | Chloramphenicol |
| NWKp027 | pBAD33t-PduA-R79E-FLAG | p15A | Chloramphenicol |
| NWKp028 | pBAD33t-PduA-R79K-FLAG | p15A | Chloramphenicol |
| NWKp029 | pBAD33t-PduA-R79Q-FLAG | p15A | Chloramphenicol |
| NWKp030 | pBAD33t-PduA-R79N-FLAG | p15A | Chloramphenicol |
| NWKp031 | pBAD33t-PduA-R79M-FLAG | p15A | Chloramphenicol |
| NWKp032 | pBAD33t-PduA-R79I-FLAG | p15A | Chloramphenicol |
| NWKp033 | pBAD33t-PduA-R79L-FLAG | p15A | Chloramphenicol |
| NWKp034 | pBAD33t-PduA-R79H-FLAG | p15A | Chloramphenicol |
| NWKp035 | pBAD33t-PduA-R79F-FLAG | p15A | Chloramphenicol |
| NWKp036 | pBAD33t-PduA-R79Y-FLAG | p15A | Chloramphenicol |
| NWKp037 | pBAD33t-PduA-R79W-FLAG | p15A | Chloramphenicol |
| NWKp038 | pBAD33t-PduA-R79G-FLAG | p15A | Chloramphenicol |
| NWKp039 | pBAD33t-PduA-R79P-FLAG | p15A | Chloramphenicol |
| NWKp040 | pBAD33t-PduA-R79*-FLAG | p15A | Chloramphenicol |

**Table S5.** Parameters used in the thermodynamic model

| | Description | Value |
|---|---|---|
| $a$ | Monomer size | $3nm$ |
| $r_c$ | Cargo radius | $2.5nm$ |
| $Q$ | Monomer number of complete shells | 32 |
| $k_b$ | bending rigidity | $50k_BT$ |
| $\theta_0$ | Preferred angle | $0.41\pi$ |
| $\epsilon_c$ | Cargo-monomer interaction | $4k_BT$ |
| $\mu_0$ | monomer chemical potential | $-6.1k_BT$ |
| $\mu_c$ | Cargo chemical potential | $-5.4k_BT$ |

**Legends for Supplementary Videos**

SI video 1:

Constant pressure, temperature all-atom simulation of native 2 PduA hexamer proteins (pdb id 3ngk) in an aqueous solution with 100 mM NaCl and 4mM $MgCl_2$. Total simulation time is 200 ns, the first 100 ns is shown in this video. The two PduA hexamers reach a stable relative orientation after about 10 ns with the help of hydrogen bonding.

SI video 2:

A coarse-grained system of 500 BMC-H, 250 BMC-T, 125 BMC-P and 250 Cargo quickly assemble into polyhedral MCPs, followed by slow relaxation of correcting  malformed units such as flipped BMC-T or BMC-P with incorrect coordination number. The interaction parameters for this simulation are the same as Fig. 2B, listed in Table S2.
